# Connectomic Organization of the Suprachiasmatic Nucleus

**DOI:** 10.1101/2024.10.20.619252

**Authors:** Jing Liu, Jing Yu, Lijun Shen, Xi Chen, Lei Ma, Hao Zhai, Linlin Li, Lina Zhang, Lu Wang, Jingbin Yuan, Bohao Chen, Bei Hong, Jiazheng Liu, Yanan Lv, Yu Cai, Yanchao Zhang, Sheng Chang, Jinyue Guo, Haoran Chen, Mianzhi Liu, Zichen Wang, Tong Xin, Fangxu Zhou, Xinghui Zhao, Dandan Zhang, Qiwei Xie, Hongtu Ma, Ying Xu, Heping Cheng, Hua Han

## Abstract

The suprachiasmatic nucleus (SCN), the mammalian master clock, is a special structure dedicated to the time computation. However, a connectomic understanding of the nucleus as a whole is lacking. Using serial section electron microscopy, we reconstructed multiscale and multimodal SCN communication networks. Intra-SCN synaptic network consists of 9,566 morphologically similar neurons and 4.3 million synapses, organized into multiscale circuitries that interact with SCN-traversing axon fascicles. Strikingly, SCN neurons engage in unique soma-soma ephaptic interaction, forming 2,038 electrotonic integrative units with the largest overlapping the light-responsive area (LRA). SCN’s paracrine modality contains 47,396 μm^3^ dense core vesicles, with notable scarcity in the LRA, suggesting cross-modal coordination and functional integration. These distinct network features provide comprehensive insights into the system architecture of the mammalian circadian clock.

## Introduction

The hypothalamic suprachiasmatic nucleus (SCN) is the central clock of circadian rhythms in mammals. It is a bilaterally symmetrical oval structure, each side containing approximately 10,000 heterogeneous neurons and hundreds of neuropeptides within a compact volume of just 0.1 mm^3^. Functionally, the SCN is synchronized to the 24-hour light-dark cycle through environmental cues and orchestrates multiscale circadian oscillations that govern physiological functions and daily behaviors(*1*). The structural basis of time information processing in the SCN differs fundamentally from that of cortical functions, such as motor, cognitive, and sensory processing(*2–7*), which have been extensively studied through connectomics. Emerging evidence suggests that the SCN employs multimodal networks of synaptic connections, neuropeptidergic signaling, and soma-soma ephaptic coupling for time computation(*1, 8*). Recent advancements in our understanding of the SCN structure include the discovery of dendro-dendritic chemical synapses(*9*) and soma-soma plate-like contact sites(*8*), as well as synaptic connections between melanopsin-expressing retinal ganglion cells and SCN neurons(*9, 10*). However, these findings are confined to partial regions of the SCN, and a global connectomic reconstruction, along with a comprehensive ultrastructural understanding of the SCN, remains an unmet challenge.

Leveraging three high-throughput single-beam scanning electron microscopes (SEM), custom computational alignment, and an automated segmentation pipeline, we have generated the first complete serial section electron microscopy (ssEM) dataset of a unilateral mouse SCN, which we have named SCNEM. This dataset offers synapse-level resolution of 5 nm × 5 nm × 40 nm and encompasses a total volume of ∼0.06 mm^3^, amounting to 88.3 terabytes. The SCNEM connectomic reconstruction provides a comprehensive perspective on communication networks within the SCN, including synaptic connectivity, ephaptic interaction networks, and neurosecretory organization. Trans-modal analysis reveals coordination among these communication modalities.

To facilitate the analysis of the mammalian circadian clock connectome, we are making the entire SCNEM publicly accessible, along with the custom-built toolset for reconstruction and interactive visualization.

## Results

### Multiscale reconstructions of the SCNEM dataset

1 mm^3^ EM tissue block containing the whole SCN was prepared from an 8-week-old male C57BL/6J mouse, which had been entrained to 12:12 light-dark cycles for two weeks before being sacrificed at Zeitgeber Time (ZT) 11.5. The tissue block was horizontally cut into serial sections with an average thickness of 40 nm, which were then collected onto tapes. Unilateral regions of interest (ROIs) within 6,824 sections were subsequently imaged at a resolution of 5 nm × 5 nm using three high throughput single-beam SEMs. Stitching and alignment yielded a ssEM volume of 384 μm × 704 μm × 273 μm (mediolateral, anteroposterior, ventrodorsal) with 88.3 terabyte voxels, which we have named SCNEM (Fig. 1, A and B). The unilateral SCN spans a volume of ∼0.06 mm^3^, with dimensions of 367 μm × 612 μm × 265 μm (Fig. 1B and supplementary materials), consistent with previous works(*11, 12*). Based on the unique appearance of densely packed neurons in SCNEM, we customized automated segmentation pipelines, particularly those for dense reconstruction, to reduce merge errors (supplementary materials). The pipelines were then used to generate a panoramic and multiscale view of SCNEM, as well as two conduit systems—capillaries and axon fascicles within the SCN (fig. S1A).

**Fig. 1.**
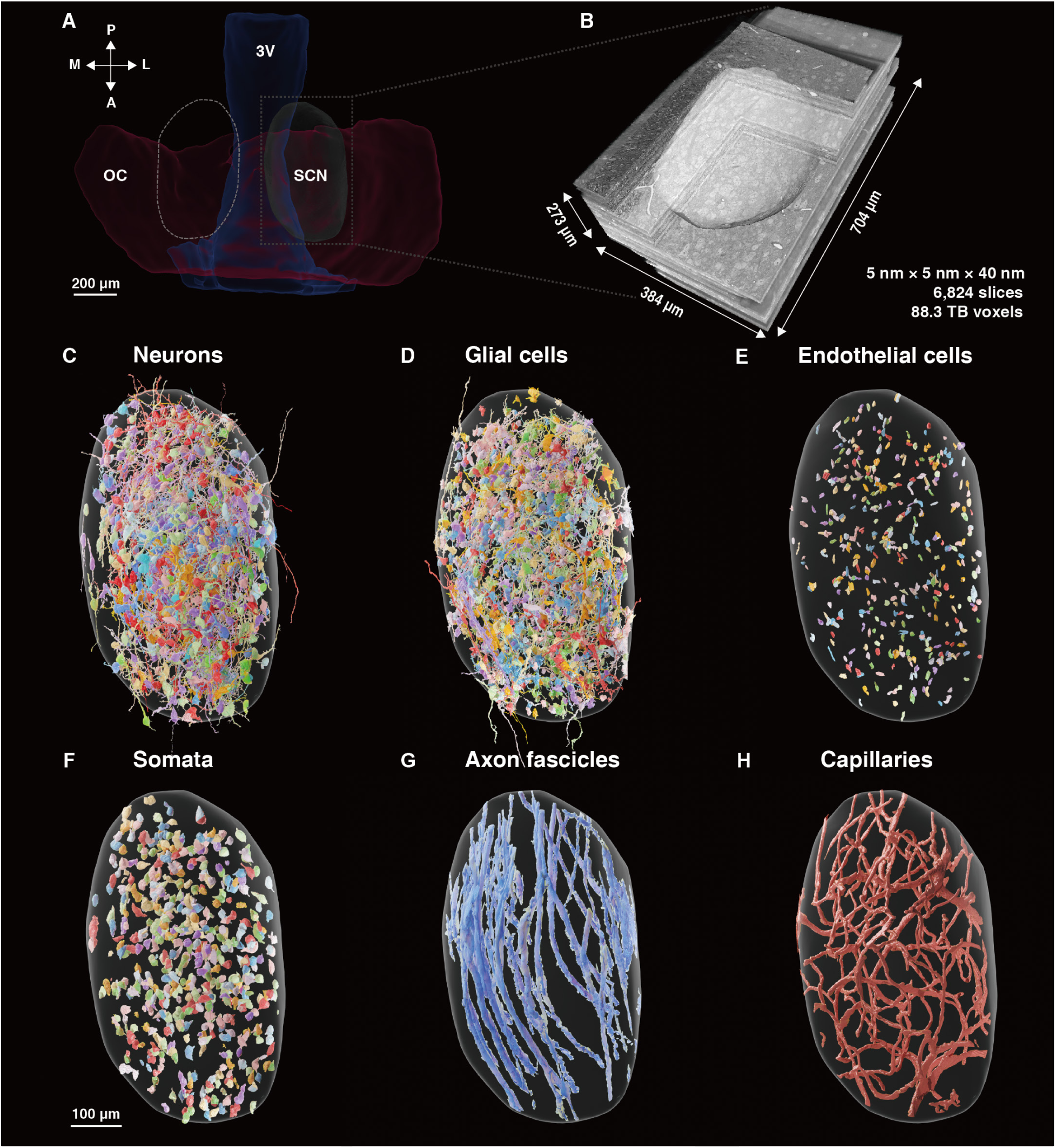
A connectomic reconstruction of mouse unilateral SCN. (**A**) 3D renderings of SCN, 3V (third ventricle) and OC (optic chiasm) estimated from micro-CT dataset in a horizontal view. Gray dashed rectangle demarcates the area for ssEM imaging. Dashed ellipsoid represents the symmetrical SCN on the opposite side. M, medial; L, lateral; P, posterior; A, anterior. (**B**) SCNEM dataset of the intact unilateral SCN, with dimensions of 384 μm × 704 μm × 273 μm. 6,824 slices were imaged at a resolution of 5 nm × 5 nm × 40 nm, with a total voxel of 88.3 terabytes. (**C** to **E**) Automated reconstructions of 9,566 neurons, 1,522 glial cells and 435 endothelial cells (shown by nuclei) in the SCNEM. Among them, 600 neurons, 594 glial cells and all endothelial cells were used for presentation. (**F**) Automated reconstructions of 11,523 somata in the SCNEM, 600 of which were used for presentation. (**G**) Automated reconstructions of 18 axon fascicles (16.1 mm in the unilateral SCN). (**H**) Automated reconstructions of capillaries (25.2 mm in the unilateral SCN). Scale bar, 100 μm.

By dense reconstruction, we identified a total of 11,523 cells, comprising 9,566 neurons (83.0%), 1,522 glial cells (13.2%) and 435 endothelial cells (3.8%) (Fig. 1, C to F and supplementary materials). The ratio of glial to neuronal cells is approximately 1:6, which is lower than previously described by single-cell analysis of gene expression(*11*). This ratio may be underestimated due to the limited understanding of glial cell morphologies in the SCN. Subcellular architectures within the SCN were also autodetected, including 4.3 million synapses, 16 million mitochondria, 47,396 μm^3^ dense core vesicles (DCVs), and 914,103 μm^3^ small synaptic vesicles (SSVs) (fig. S1). Furthermore, we identified distinctive cable-like structures—axon fascicles— totaling 16.1 mm within the SCN, each comprising up to 1,200 axons and traversing the SCN along the anteroposterior axis (Fig. 1G). Meanwhile, we reconstructed the global architecture of 25.2 mm microvasculature across the SCN (Fig. 1H and fig. S1B, Capillaries). This intricate network traverses the nucleus, featuring a branch point density of 6.6 × 10^-6^ per μm^3^ and vessel radii ranging up to 6.5 μm, consistent with previous reports(*13*). A comprehensive view of these global features in an intact unilateral SCN is presented in movie S1.

### The synapse-resolution connectome of the SCN

A synapse-resolution connectome of the SCN is quintessential for comprehending the structural and functional principles of time computation by the master clock. After automatic agglomerations, 391 neurons were proofread, compartmentalized, and assigned with detected synapses (Fig. 2C, table S1 and movie S2). These neurons were categorized into a single class with three subtypes based on similarities in gross morphology and variations in dendritic arborization patterns. The subtypes, distinguished by the number of dendrites radiating from the cell body(*12*), include uni-dendritic neurons (UNs), bi-dendritic neurons (BNs), and multi-dendritic neurons (MNs). Their relative abundances were 9.5%, 72.6% and 17.9% for UNs, BNs and MNs, respectively, and different subtypes entwine irregularly to form a mosaic pattern within the SCN (Fig. 2, A and C and movie S2).

**Fig. 2.**
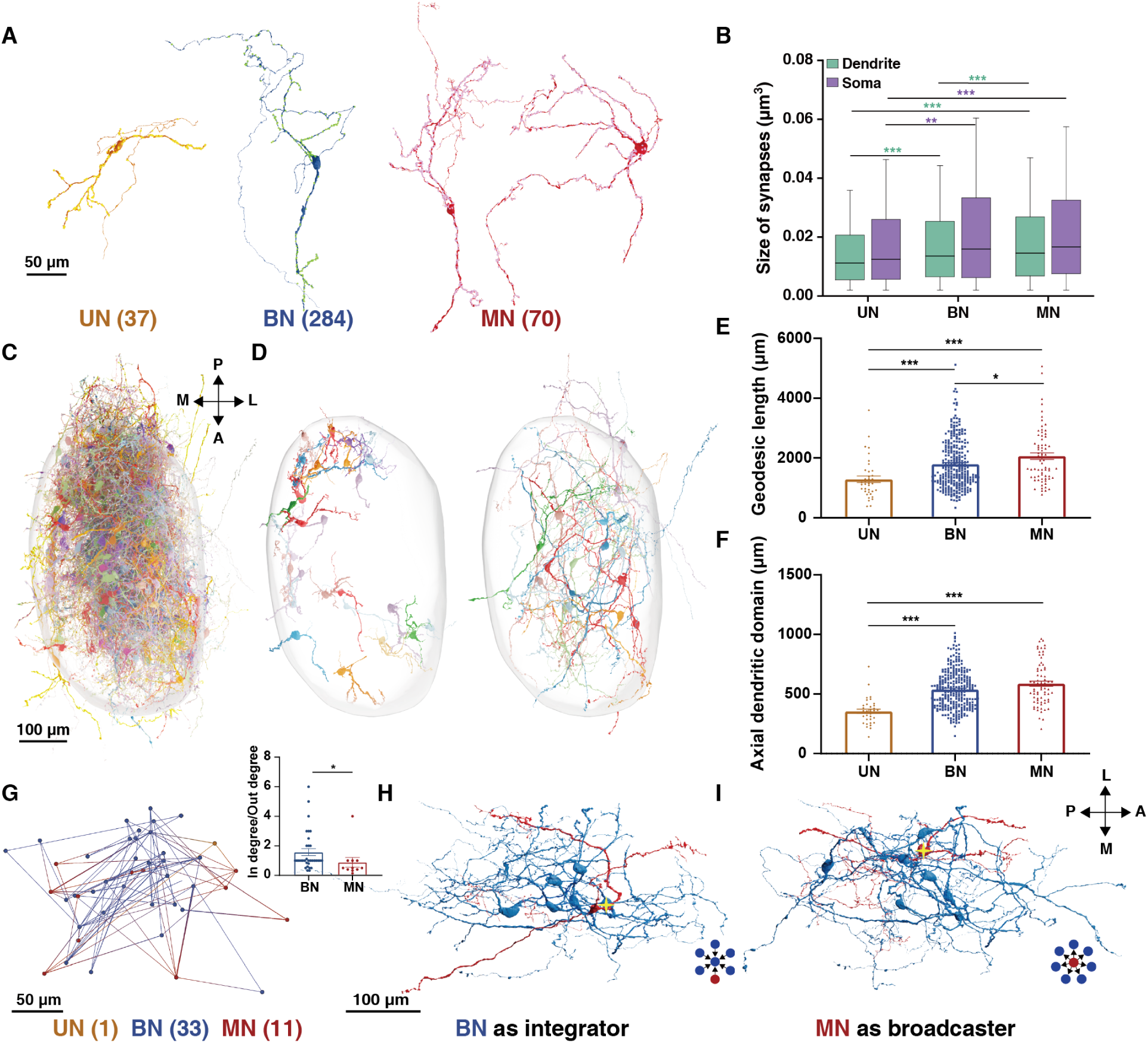
Synaptic connectivity network in the SCN. (**A**) Representative morphology of neurons in UN (yellow), BN (blue), MN (red) subtypes. Scale bar, 50 μm. (**B**) Size of synapses located on dendrites and somata. Black lines denote the medians. For synapses on dendrites, n = 8,572 (UN), 106,756 (BN) and 29,040 (MN). For synapses on somata, n = 522 (UN), 5,092 (BN) and 1,312 (MN). (**C**) 3D renderings of 391 proofread neurons in the SCN (See also movie S2). Scale bar, 100 μm. (**D**) Intra-SCN neuronal dendritic projection. Out of 42 neurons, 26 neurons with short-range axial dendritic domains (<300 μm, left) and 16 neurons with long-range axial dendritic domains (>950 μm, right) were shown. (**E** and **F**) Geodesic length and axial dendritic domain. Data are shown as mean ± s.e.m.. n = 37 (UN), 284 (BN) and 70 (MN). (**G**) The largest strongly connected component (45 nodes and 107 connections) identified in the synaptic connectivity graph of 391 proofread neurons. Color code: UN node (yellow), BN node (blue), MN node (red). Inset: the ratio of in-degree to out-degree in BN and MN subtypes. Data are shown as mean ± s.e.m.. n= 33 (BN) and 11 (MN). Scale bar, 50 μm. (**H**) A BN serves as an integrator neuron (cross star), receiving 6 connections from other neurons via 6 synapses. Inset: sketch of connectivity pattern. Color code: BN (blue), MN (red). (**I**) A MN serves as a broadcaster neuron (cross star), sending 7 connections to other neurons via 8 synapses. Inset: sketch of connectivity pattern. BN (blue), MN (red). Scale bar, 100 μm. M, medial; L, lateral; P, posterior; A, anterior.

The automated assignment of synapses to the proofread neurons revealed diverse synaptic configurations. Axo-dendritic and axo-somatic synapses constitute nearly half of all synapses (51.6%), while dendro-dendritic synapses, dendro-somatic, and somato-dendritic (47.5%) facilitate direct communication between two dendrites or between a dendrite and a soma, which can be serial or reciprocal(*14*)^,16^ (fig. S2A, top). Additionally, we identified a previously undescribed rare type of synapse in large-scale EM reconstruction, the axo-axonal synapse, accounting for 0.8% of the total (fig. S2A, bottom). Given that synapse size correlates with physiological strength(*15, 16*), we showed that synapses located on dendrites and somata were smaller than those on axons, suggesting distinct functional roles for different neuronal compartments (fig. S2B; dendrites versus axons: *p* < 0.001; somata versus axons: *p* < 0.001; Mann–Whitney *U*-test). Synapses located on dendrites and somata, which primarily constitute neuronal inputs, were smallest in UNs, followed by BNs and MNs (Fig. 2B; UN versus BN: *p* < 0.001 (dendrites), *p* < 0.01 (somata); UN versus MN: *p* < 0.001 (dendrites), *p* < 0.001 (somata); Mann–Whitney *U*-test). The number of synapses per neuron, excluding those on the axons, ranges from 51 to 1,202, with an average of 387. By neuronal subtypes, the synapse counts average 246 for UNs, 394 for BNs and 434 for MNs. To mitigate the influence of neuronal length and surface area, we further examined the linear or areal synapse density, defined as the number of synapses per 1 μm of path length or per 1 μm^2^ of surface area, respectively. The linear synapse density on dendrites and areal synapse density on somata, averaging 0.2 synapse/μm and 0.02 synapse/μm^2^ per neuron, were found higher in MNs and BNs compared to UNs (fig. S2C; UN versus BN: *p* < 0.05 (dendrites), *p* = 0.0881 (somata); UN versus MN: *p* = 0.1749 (dendrites), *p* = 0.0823 (somata); Mann–Whitney *U*-test). Thus, although the SCN is rich in synaptic types, its synapse density, whether per neuron or per unit of path length, is significantly lower than those found in the cortex and hippocampus. (e.g., >10,000 synapses and ∼1.5 synapses per 1 μm per giant pyramidal neuron)(*17–19*).

SCNEM enables us to quantify the projection of complete dendritic arbors, another key determinant of intra-SCN synaptic connectivity and communication. To assess a neuron’s receptive field, we calculated the geodesic length and axial projection distance of its dendritic arbors, and defined the axial dendritic domain as the sum of axial projection distances along the mediolateral, anteroposterior and ventrodorsal axes (see supplementary materials). We found that both the path length and axial dendritic domain usually followed the order of MN > BN > UN (Fig. 2, E and F; *p* < 0.001 for all (geodesic length); MN versus UN: *p* < 0.001, BN versus UN: *p* < 0.001, MN versus BN: *p* = 0.0796 (axial dendritic domain); Mann–Whitney *U*-test; see also fig. S2, D to F). Among the 391 proofread neurons, we identified 16 long-range neurons with axial dendritic domains exceeding 950 μm, and 26 short-range neurons with axial dendritic domains less than 300 μm (Fig. 2D). The long-range neurons can thus span almost the entire SCN, covering 70–86% of each axial direction; while even the short-range ones can integrate inputs from a substantial portion of the nucleus, covering 19–34% of the axial dimensions (Fig. 2D and movie S3). This extensive dendritic arborization affords the basis for the formation of multiscale micro-circuitries within the clock. In particular, BNs and MNs, with larger synapse size, higher synaptic density, and larger axial dendritic domains, likely play a more significant role in synaptic communication within the SCN.

By linking synapses to their pre-synaptic and post-synaptic neurons, we identified 300 connections and constructed a synaptic connectivity graph among the 391 neurons (movie S3). Although this connectivity analysis covers only 4.1% of the entire neuronal population, a large strongly connected component (SCC) consisting of 45 neurons was identified (Fig. 2G). This SCC corresponds to a connection probability of 0.054 and an average clustering coefficient of 0.11, both comparable to those found in the cortex(*16*). The average shortest path length in this SCC is 3.99, indicating that any two nodes can typically reach each other within 4 steps, suggesting highly efficient communication. We found that MNs were more likely to function as broadcasters, while BNs served as integrators (Fig. 2G, inset; *p* < 0.05, Mann–Whitney *U*-test). In particular, we pinpointed a BN that acts as an integrator (ratio of in degree and out degree = 6:1), receiving signals from 6 neurons via 6 synapses, and an MN that acts as a broadcaster (ratio of in degree and out degree = 1:7), sending signals to 7 neurons via 8 synapses (Fig. 2, H and I).

Overall, the neuronal architecture underlying SCN time information processing is characterized by a uniform neuronal type, low synaptic density, diverse synaptic types, extensive dendritic projections, and highly efficient connectivity, leading to multiscale circuitries. These structural features collectively support the notion that the SCN functions as an indivisible entity in the genesis of complex physiological rhythms.

### Axon fascicles traversing the SCN

A distinctive feature of the SCN is the grouping of axons into 18 cable-like axon fascicles. These unique structures prompt numerous questions about the nature of their information flow, origins, and destinations. Through automated segmentation followed by manual proofreading, we showed that nearly all of them traverse the nucleus along the anteroposterior axis and often disperse into smaller fascicles (Fig. 3A). Composed primarily of unmyelinated axons(*12*), these fascicles appear to originate from and terminate in regions outside the SCN, but their trajectories differ from that of myelinated axons from the optic chiasm, which primarily extend along the mediolateral axis (fig. S3, A and B). This suggests that the axon fascicles may function as conduits connecting different hypothalamic regions rather than receiving photonic signals from the retina. Incorporating dense reconstructions at 10 μm intervals along the fascicle skeleton (Fig. 3, B to D and supplementary materials), we found that each axon fascicle contains 99 to 1,191 axons, with the number of axons linearly correlated with the fascicle’s cross-section area, which ranged from 6.3 to 89.1 μm^2^ (Fig. 3E, *r* = 0.9641, *p* < 0.0001, Pearson correlation analysis).

**Fig. 3.**
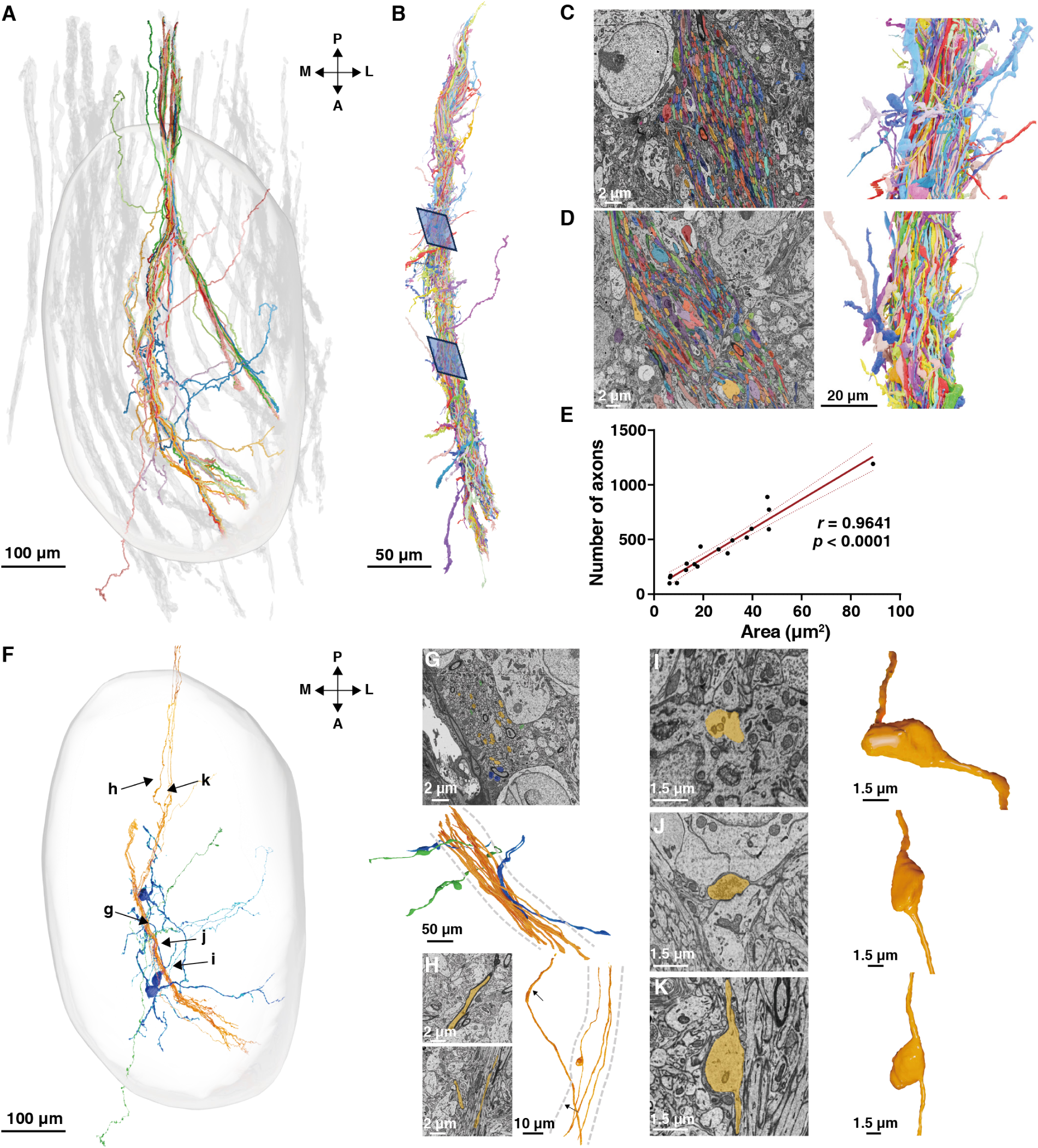
Axon fascicles in the SCN. **(A**) 3D renderings of axon fascicles identified in the SCNEM. Individual fascicles contain approximately 100–1,200 axons. 75 randomly selected axons within and around a representative fascicle were highlighted by color. Scale bar, 100 μm. (**B**) A representative single axon fascicle with all assigned axonal fragments. Blue rectangles indicate two ROIs shown in panels (C and D). Scale bar, 50 μm. (**C** and **D**) EM images (left) and 3D renderings (right) of two ROIs in (B). Scale bars, 2 μm and 20 μm. (**E**) Linear correlation between the cross section area and the number of axons within fascicles. Red line represents the fitted slope, and dashed lines indicate the 95% confidence band. The correlation coefficient *r* and *p* value were obtained from a two-tailed, non-parametric Pearson correlation analysis. n = 18. (**F**) Trajectories and orientations of 23 proofread axons. Color coding of orientation: orange, axons that traverse the SCN along the fascicle without leaving; blue, axons originating from SCN neurons and merging into the fascicle; and green, axons crossing the fascicle. Black arrows indicate the positions shown in panels (G to K). Scale bar, 100 μm. (**G** and **H**) Magnified views corresponding to the positions in (F). Note the different orientations of axons in the fascicle. Gray dashed lines mark the boundary of fascicle. Scale bars, 2 μm, 10 μm and 50 μm. (**I** and **J**) EM images (left) and 3D renderings (right) of synapses formed between axon fascicles and SCN neurons, axon fascicles as presynaptic (J) or postsynaptic (I) structures. Scale bars, 1.5 μm. (**K**) EM images (left) and 3D renderings (right) of an axonal bouton in the fascicle. Scale bars, 1.5 μm. Renderings of (G to K) are smoothed using local alignment. M, medial; L, lateral; P, posterior; A, anterior.

To explore axon-fascicle interactions within the SCN, we manually traced the skeletons of 75 randomly selected axons within and around a fascicle’s contour (Fig. 3F). We found that 68 axons (90.7%) passed through the SCN without leaving the fascicle, 3 axons (4%) ran through the fascicle perpendicularly, and 3 axons (4%) originating from SCN neurons merged into the fascicle briefly (Fig. 3G). Additionally, 1 myelinated axon (1.3%) exited the fascicle and entered the SCN (Fig. 3H). Detailed examination of 23 axons revealed bouton-like structures extending from the fascicle, forming synapses with SCN neurons at various positions (Fig. 3, I to K). These findings suggest that axon fascicles may serve as a novel information highway, connecting surrounding hypothalamus nuclei with sparse entry and exit points along their route through the SCN, thereby integrating SCN time information with other hypothalamic functions.

### Ephaptic interaction network in the SCN

Soma-soma ephaptic interaction, characterized by large plate-like contacts between clustered neurons, is a feature that has not yet been reported in any other brain regions(*8*). Although the exact functional role of this interaction remains unclear, it is thought to serve as an electrophysiological substrate for synchronization, potentially contributing to the maintenance of circadian rhythms(*8, 12*). Theoretically, the spatial spread of electrotonic field potential extends farther when extracellular electrical resistivity is increased by reducing aqueous extracellular space of high conductivity, rendering a greater electrical space constant. Therefore, we refer to a neuronal cluster coupled through ephaptic interactions as an electrotonic integrative unit (EIU).

Using SCNEM, we constructed a weighted adjacency graph of all SCN neurons, comprising 9,566 nodes and 11,485 edges (Fig. 4A). In this graph, each node represents a soma, and each edge reflects the strength of ephaptic interaction between adjacent somata, quantified by their contact surface area. Applying the minimum cut algorithm (supplementary materials) decomposed the graph into a complete set of EIUs, revealing 2,038 EIUs of varying sizes and 10,169 edges with an average contact area of 7.4 μm^2^ (Fig. 4, B and C and fig. S4A). These contacts together cover an average of 5.4% and a maximum of 45.4% of the soma area (Fig. 4D). The node degree, defined as the number of contacting somata, ranges from 0 to 12 and closely follows a Poisson distribution (Fig. 4E, R^2^ = 0.9067). Strongly connected nodes, characterized by a node degree greater than five, are predominantly located in the ventral and anterior regions of the SCN (fig. S4B). In contrast, isolated somata (0 node degree) account for ∼12.6% (1,203 out of 9,566) of the neuronal population and are uniformly distributed across the SCN (fig. S4C).

**Fig. 4.**
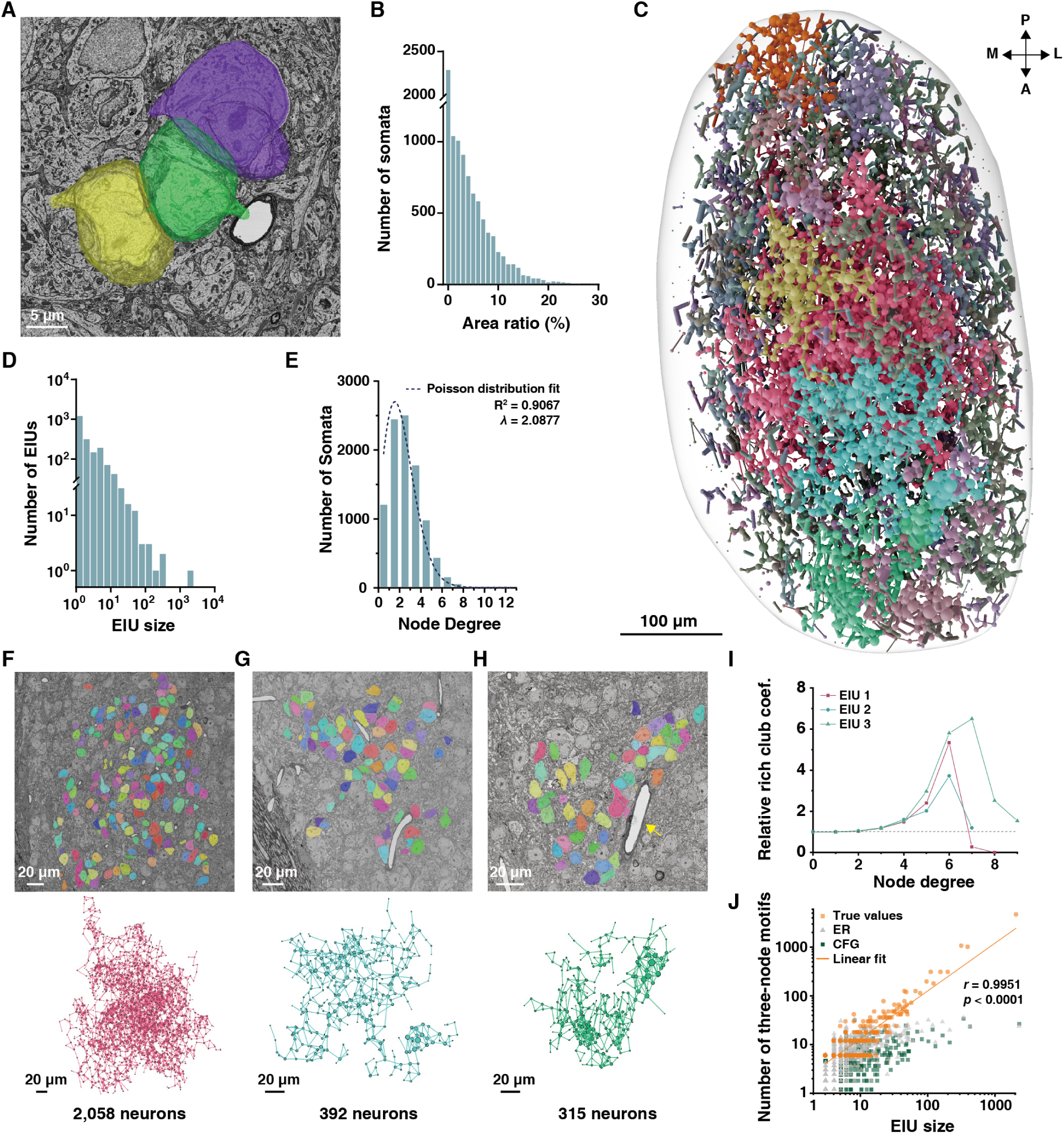
Soma-soma ephaptic interaction network in the SCN. (**A**) Representative soma-soma contact sites among three somata. Scale bar, 5 μm. (**B**) EIU size among all EIUs. n = 2,038 EIUs. (**C**) A comprehensive soma-soma ephaptic interaction network consisting of 9,566 SCN neurons. 2,038 EIUs in total, each represented by a different color (See also movie S3). Each node represents a soma and each edge represents the contact between two adjacent somata. Scale bar, 100 μm. M, medial; L, lateral; P, posterior; A, anterior. (**D**) Distribution of the ratios of total contact area to soma surface area. n = 9,566 neurons. 15 observations with ratios greater than 30% are omitted from the plot. (**E**) Node degree among all EIUs. R^2^ refers to the fitting coefficient and *λ* represents the expected node degree in the best-fit Poisson distribution. n = 9,566 neurons. (**F** to **H**) Representative EM images (top) and network connectivity graph (bottom) of three largest EIUs. Node size reflects the node degree of the respective soma. The yellow arrow indicates a capillary adjacent to the chain-like EIU. Scale bars, 20 μm. (**I**) Relative rich club coefficient as a function of node degree for three EIUs, computed relative to 1,000 randomly sampled CFG models. Relative rich club coefficients exceeding 1 (black dashed line) indicate the presence of a rich club architecture. Notably, when node degree reaches 7 in EIU1, the network becomes sparser compared to CFG model. (**J**) Comparison of three-node motif frequency against EIU size. The true data is represented by orange circles, the ER model by grey triangles, and the CFG by green squares. Orange line represents a linear fit to the observed data. The correlation coefficient *r* and *p* value were obtained from a two-tailed, non-parametric Pearson correlation analysis. n = 2,038 EIUs.

Strikingly, we identified three large EIUs comprising 2,058, 392 and 315 neurons, respectively, located in the central, ventral, and anterior regions of the SCN (Fig. 4 F to H, and movie S4). The largest EIU features a spherical structure that grossly overlaps the light-responsive area (LRA)(*20, 21*) recently identified in the core region of SCN (Fig. 4F). The second largest EIU primarily extends along the anteroposterior axis, forming a planar structure, while the third largest displays a distinct chain-like structure in the anterior region(*12*) (Fig. 4H). Plots of the relative rich club coefficient as a function of node degree, computed relative to configuration graph (CFG)(*22*), reveal abundant hub nodes (node degree greater than 3) in soma-soma ephaptic interactions across all three large EIUs (Fig. 4I and supplementary materials), which help to integrate and distribute information due to their high connectivity.

The abundance of three-node motifs, the smallest circuits within a network, reflects a fundamental network property. An over-representation of these motifs compared to random graphs is a typical characteristic of cortical synaptic wiring networks(*17*). Visual inspection suggests a similar over-representation in the SCN ephaptic interaction network. Analyses using both Erdős– Rényi (ER) graph and CFG(*22*) confirmed this phenomenon, particularly in larger EIUs (Fig. 4J). Additionally, strong linear correlation between EIU size and three-node motif count (Fig. 4J, *r* = 0.9951, *p* < 0.0001, Pearson correlation analysis) underscores another inherent property of this ephaptic interaction network throughout the SCN. Overall, the extensive soma-soma ephaptic interactions may provide a unique modality of signal integration and synchronization in the SCN.

### Neurosecretion organization in the SCN

Neurons in the endocrine hypothalamus, including those in the SCN, are highly active in releasing neuropeptides like arginine vasopressin (AVP), vasoactive intestinal polypeptide (VIP), and cholecystokinin (CCK)(*23–26*). This neurosecretion is organized into two subsystems, fast-acting synaptic neurotransmitter release from SSVs and slower modulatory neuropeptide release from DCVs(*1, 27*). To examine SSV and DCV distribution, an automated method was applied to identify regions of DCV and SSV accumulation (Fig. 5A and supplementary materials). Our results showed that SSVs and DCVs within the SCN occupied 914,103 μm³ and 47,396 μm³, respectively, representing 95.1% and 4.9% of the total vesicle volume. Regarding vesicle partitioning among neuronal compartments, we found that vesicle docking predominantly occurred in neuronal dendrites or axons, with somatic SSVs and DCVs accounting for only 0.4% and 1.4% of their respective total volumes. We further investigated somatic vesicle distribution across neuronal subtypes and found that SSV volumes were comparable among them, while UNs exhibited a higher volume of DCVs, suggesting a more prominent role for UNs in paracrine neuropeptidergic signaling (Fig. 5, F to H).

**Fig. 5.**
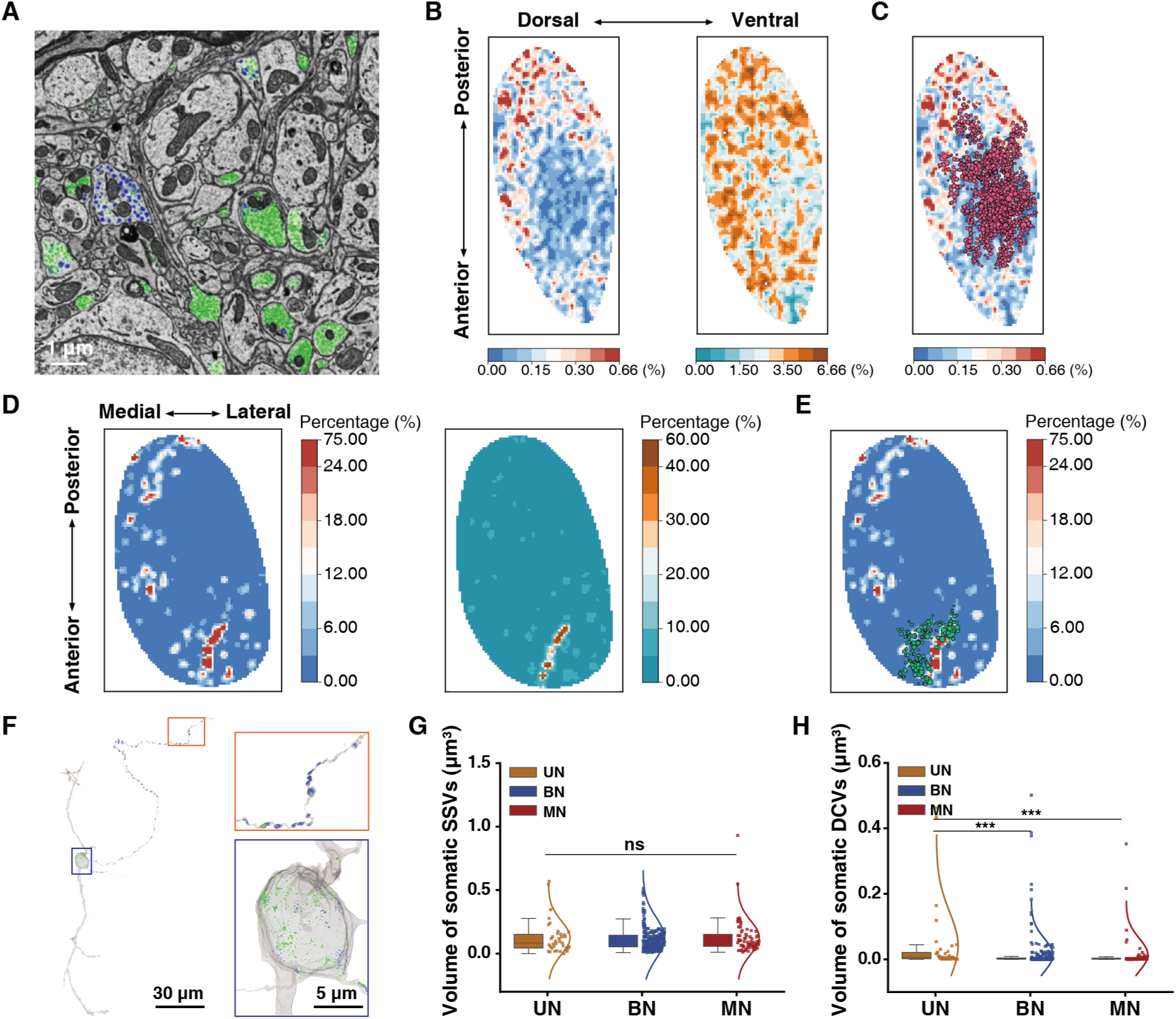
Neurosecretory organization in the SCN. (**A**) Vesicle classification. Green represents SSVs and blue represents DCVs. Scale bar, 1 μm. (**B**) Maps of DCV (left) and SSV (right) densities in the 350-μm sagittal section. The density is denoted by vesicle volume fraction within a cube of 10 × 10 × 10 μm^3^. (**C**) Map of DCV density (left panel in (B)) overlaid with the largest EIU consisting of 2,058 neurons. (**D**) Maps of somatic DCV (left) and SSV (right) densities in the 70-μm horizontal section. The density is denoted by the ratio between the intra-soma value and the overall value within a cube of 10 × 10 × 10 μm^3^. (**E**) Map of somatic DCV density (left panel in (D)) overlaid with the third chain-like EIU. (**F**) Distribution of DCVs and SSVs in a representative neuron. Inset shows magnified views of the soma (blue squares) and axon (orange squares). Scale bars, 30 μm and 5 μm. (**G** to **H**) Somatic SSV (G) and DCV volumes (H) in UN, BN, and MN subtypes. n = 37 (UN), 284 (BN) and 70 (MN). Black lines denote the medians.

The density maps of SSVs show a relatively uniform distribution with only minor fluctuations across the nucleus (Fig. 5B, right). In contrast, DCVs show a higher volume fraction in the peripheral SCN, with a notable scarcity in the central SCN (Fig. 5B, left). This central scarcity coincides with the largest EIU of the ephaptic interaction network (Fig. 5C). In somatic vesicle density maps, a distinct chain-like substructure is evident in the anterior SCN, characterized by high somatic densities of both DCVs and SSVs (Fig. 5D), corresponding to the chain-like, third largest EIU (Fig. 5E). It is noteworthy that these chain-like substructures are adjacent to capillaries (Fig. 4H, top), suggesting a paracrine module for long-range signaling via blood vessels.

### Transmodal coordination

The results above show that the reconstruction of multimodal networks unexpectedly uncovered the alignment between two largest EIUs and the neurosecretory substructures. Motivated by this finding, we systematically investigated the spatial architectures of all reconstructed attributes across the entire SCN in three dimension (3D), planar projection (2D), and axial projection (fig. S1 and supplementary materials). As illustrated in fig. S1, B to D, spatial density fluctuations are a common feature shared by all examined attributes, leading to the emergence of intermediate-scale substructures on 2D and 3D maps, as well as global gradients and local minima and maxima along the SCN axes. For instance, in addition to the density fluctuations of DCVs and SSVs, soma density exhibits a dorsal-ventral gradient, with higher density but smaller soma size in the ventral region (fig. S1, B to D, somata), consistent with the functional modular organization of the SCN(*28*).

To quantify transmodal coordination, we conducted Pearson correlation analyses of spatial density fluctuations between pairs of attributes (fig. S5A and supplementary materials). As expected, synapses exhibited the highest positive correlation with SSVs (see also fig. S5B, *r* = 0.7983, *p* < 0.0001, Pearson correlation analysis). Likewise, DCVs also showed a linear correlation with both synaptic density and SSVs (fig. S5C, synapses: *r* = 0.3933, *p* < 0.0001; SSVs: *r* = 0.6208, *p* < 0.0001; Pearson correlation analysis). These pairwise correlations suggest that synaptic transmission via SSVs and paracrine signaling via DCVs operate in a coordinated manner within the SCN. In contrast, both subsystems of neurosecretion correlated negatively with soma density (SSVs: *r* = -0.4391, *p* < 0.0001; DCVs: *r* = -0.3423, *p* < 0.0001; Pearson correlation analysis), such that higher soma density was associated with a lower vesicle volume fraction (see Discussion).

Regarding the metabolic modality of the SCN, we observed strong spatial coordination between mitochondrial and synaptic densities (*r* = 0.6960, *p* < 0.0001, Pearson correlation analysis), as well as between mitochondria and SSVs and DCVs (fig. S5A, SSVs: *r* = 0.5378, *p* < 0.0001; DCVs: *r* = 0.2897, *p* < 0.0001; Pearson correlation analysis). This suggests that regions with higher synaptic and paracrine activity are supported by an elevated energy supply, and vice versa. These results underscore the essential role of mitochondrial bioenergetics and metabolism in sustaining both synaptic transmission and paracrine communication (fig. S5, D and E). Conversely, capillaries showed no nucleus-wide correlation with other modalities (somata: *r* = 0.0386; synapses: *r* = -0.0549; mitochondria: *r* = -0.0445; SSVs: *r* = -0.0614; DCVs: *r* = -0.0445; Pearson correlation analysis), except for its local coincidence with the chain-like vesicle and EIU substructures (see above). This result implies a homogeneity of basal metabolism across the SCN. Overall, our quantitative analysis reinforces the idea that the multimodal networks within the SCN interact and coordinate to operate as an integrated system.

## Discussion

The hypothalamus is a paired structure located below the thalamus in the vertebrate brains, containing several nuclei responsible for regulating essential body functions. Among these, the SCN stands out as a unique central nervous nucleus dedicated for the integration, computation and representation of circadian time information. However, a connectomic understanding of SCN is lacking. Using three high throughput single-beam SEMs combined with automated segmentation and subsequent manual proofreading, we generated the first synapse-resolution ssEM dataset of the entire mouse unilateral SCN, and conducted a comprehensive connectomic analysis. In this way, we provided multiscale and panoramic views of the multimodal communication networks within the SCN, including synaptic wiring diagrams, ephaptic interactions and neurosecretory organization, as well as their metabolic substrates (Fig. 6A). In addition, transmodal correlation analysis revealed distinct patterns of coordination among these networks across the SCN.

**Fig. 6.**
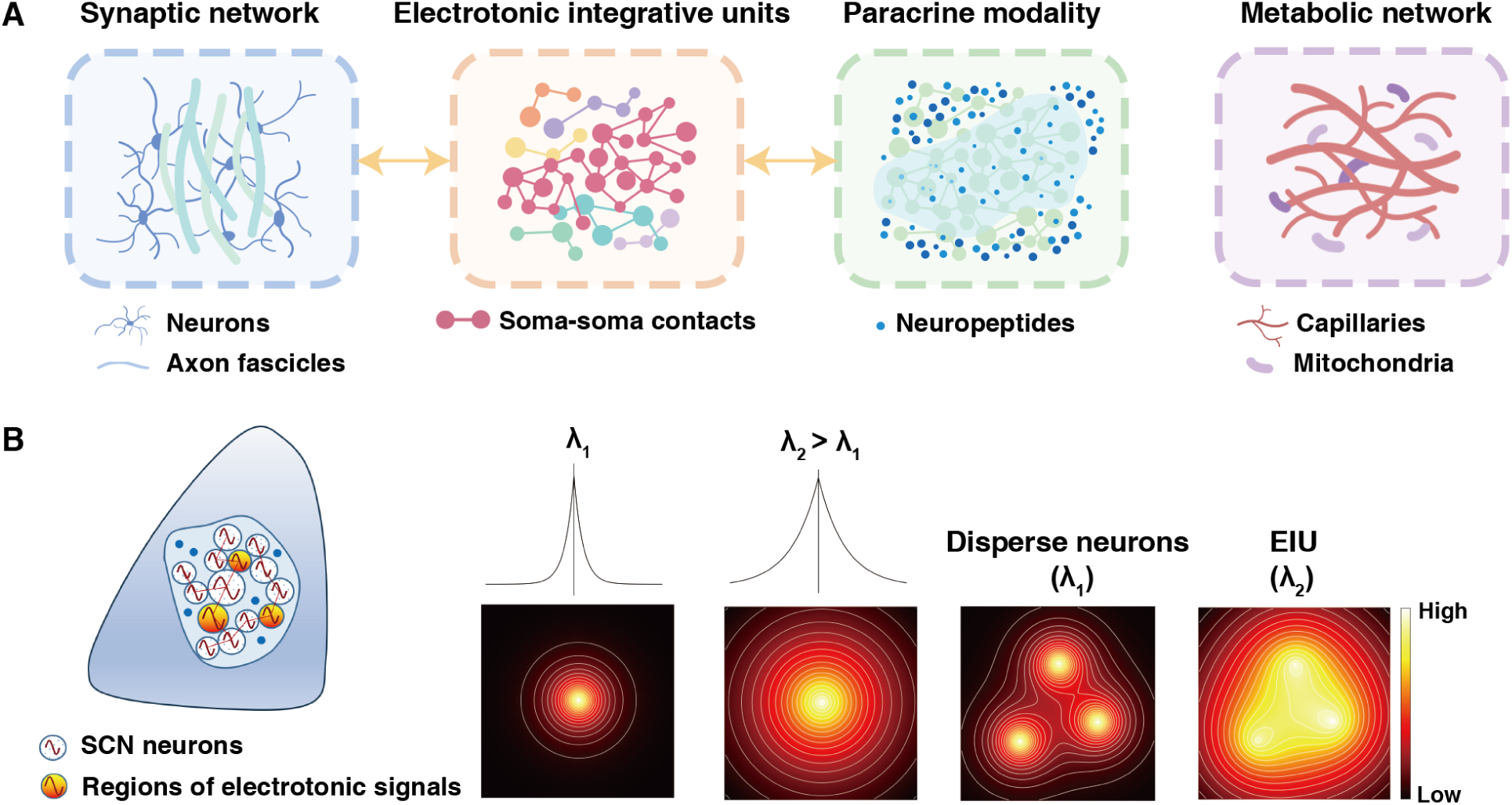
Multimodal networks in the SCN. (**A**) Schematic representation of SCN networks. From left to right: synaptic network comprising morphologically similar neurons, interacting with axon fascicles traversing the nucleus; ephaptic interaction network with 2038 EIUs; paracrine modality with a notable scarcity in the largest EIU; metabolic network containing capillaries and mitochondria. (**B**) A working model for integration of electrotonic signals in an EIU. Because of soma-soma interaction, extracellular electrical resistivity is higher compared to the situation with only disperse neurons, resulting in an increased electrical space constant (λ_2_ > λ_1_). This larger space constant enhances the spatial spread of electrotonic signals, and thus facilitates summation of signals from multiple regions in the same EIU.

Our study has uncovered salient features of the SCN synaptic network. Among the 391 proofread SCN neurons, we found no distinct morphological differences, except for variations in the number and extent of dendritic arborization. This finding suggests a system-wide coherence, indicating that the entire SCN is likely composed of morphologically similar neurons, unlike the diverse neuron types found in the cortex(*2, 3, 17*). This structural uniformity supports our recent discovery that the SCN operates as a unified entity for time computation and representation via a group decision-making mechanism, with each neuron contributing uniquely yet uniformly(*29*). Functionally, almost all SCN neurons are GABAergic, but the effect of GABA varies depending on the dynamic state of the SCN networks and the expression of chloride transporters(*30–32*). It thus remains challenging to definitively classify neurons in the SCN as excitatory or inhibitory. This complexity underscores the need for a deeper understanding of SCN synaptic dynamics to fully elucidate how this network orchestrates precise time computation across ∼20,000 heterogenous neurons.

Despite its apparent simplicity, the SCN’s time computation relies on a complex intra-SCN synaptic network, characterized by diverse synaptic types, extensive dendritic projections, and highly efficient micro-circuits. Synapse densities in the SCN (∼450 per neuron, 0.07 per μm^3^) are lower than those in the cortex (∼8,000 per neuron, 1.1 per μm^3^ in 0.003 mm^3^ mouse primary visual cortex)(*17, 18*). However, the SCN is rich in synaptic types, including a newly identified, rare axo-axonal synapse. The extensive dendritic projections enable all SCN neurons, whether short-range or long-range, to receive inputs from a large portion of the unilateral SCN, enhancing intra-SCN connectivity and computation. A representative micro-circuit of 45 neurons, identified among the 4.1% proofread population, includes both broadcaster and integrator neurons and demonstrates a low average shortest path length—key characteristics of a small-world network that facilitates efficient communication(*33, 34*). These multi-level components are fundamental in constructing multiscale micro-circuitries within the SCN connectome.

Additionally, we identified 18 prominent intra-SCN axon fascicles that both originate from and project to regions outside the SCN, likely representing long-range communication within the hypothalamus. SCN neurons occasionally form synapses with these axon bundles, suggesting that SCN time-processing functions are integrated with other hypothalamic activities, allowing for input signal integration and output signal sorting.

Ephaptic interaction, not yet reported in any other brain regions(*8*), constitutes a unique modality of inter-neuron communication within the SCN. An EIU, composed of a cohort of neurons with direct soma-soma contact, is thought to function as a module for integrating electric signals and facilitating synchronization. Extracellular electrical resistivity can affect the spatial spread of electrotonic field potential. In this scenario, a large EIU can summate weak electrotonic signals from the entire cohort, such that the integrated signal might become strong enough to influence the activity of all cohort members collectively and synchronously (Fig. 6B). This interpretation aligns with the striking fact that the largest EIU, comprising 2,058 neurons, overlaps the LRA, as if ephaptic interaction facilitates the entrainment of the time-keeping system by environmental light. It is tempting to speculate that this EIU may partake the integration of photic signals for the evening and morning oscillators(*20, 21, 35*). In addition, other sizable EIUs identified may also create local functional modules that integrate and synchronize activities of topological specificity.

Circadian oscillations are generated by mechanisms operating across different timescales, involving both rapid synaptic and ephaptic signaling, as well as relatively slow paracrine signaling. Global maps of DCVs depict the organization of paracrine signaling, the secretion of SCN neuropeptides such as AVP, VIP, and CCK. Intriguingly, DCVs are scarce in the core region of the SCN, which overlaps with the largest EIU and the LRA. This pronounced radial gradient of DCVs suggests coordination between neurosecretory and ephaptic interaction modalities. However, it is important to note that the mouse was sacrificed before lights off, so the observed organization could reflect physiological and regional fluctuations in the DCV pool at different times of the day. Further investigation is required to distinguish between these possibilities.

At intermediate scales, substructures in the form of axial or planar gradients and 3D fluctuations are common across all attributes examined. Connectomic transmodal correlation analysis of these substructures revealed three significant features. First, there is a positive correlation between pairs of synapses, SSVs, and DCVs, indicative of systematic coordination between synaptic transmission and paracrine signaling. Second, a negative correlation exists between soma and vesicle densities, indicating an inverse relationship between these two attributes. Third, mitochondrial density is positively correlated with the densities of synapses, SSVs or DCVs, but not with somata. This finding highlights the importance of bioenergetic support for both synaptic and neurosecretory activities and is consistent with the notion that ephaptic integration may require minimal energy input.

In summary, this study provides the first system-level EM analysis of a hypothalamic nucleus, revealing complex cross-scale organization features within the synaptic, EIU, and neurosecretory networks of the mammalian circadian clock. Alongside recent functional insights from large-scale Ca^2+^ imaging and machine learning(*29*), our findings support the concept that the SCN functions as a unified entity with multiscale and multimodal organization(*36*). Investigating the hypothalamic nucleus poses unique challenges compared to the cerebral cortex(*2, 3, 6, 7*) due to its deep anatomical location, bilateral symmetry, and complex array of inputs and outputs. Our work establishes a new analytical framework for studying the hypothalamic nucleus, which may inspire further structural, functional, and computational research to uncover fundamental system design principles in other hypothalamic nuclei.

## Acknowledgments

We thank Y.X. Wu, X.L. Wen and other staffs from Grand View (Beijing) Technology Co., Ltd for ground truth annotation, neuronal skeleton labeling, and proofreading; W.L. Xiao, J.M. Liu, C. Zhao and B.F. Dai for assistance with the proofreading software. Q.C. Zhai, X.H. Fang and D. Zhang for helpful discussion; Y.T. Chen for assisting with EM imaging and quality supervision. We also thank the Transdisciplinary Platform of Functional Connectome and Brain-inspired Intelligence in Huairou Science City in Beijing and the Microscopic Technology & Analysis Center at the Institute of Automation, Chinese Academy of Sciences for providing technical support and device resources, Biomedical Computing Platform of the National Biomedical Imaging Center at Peking University for providing the necessary computational resources.

## Author contributions

H.C. and H.H. conceptualized and supervised the project. J.Y. designed and performed the animal experiment. J.Y and L.L. developed the sample preparation methods and prepared the EM samples. L.L. performed the collection of serial sections. L.Z. and H.M. performed EM imaging. X.C., B.C., S.C., Y.L., H.R.C. and T.X. performed the stitching and alignment. J.L., J.G., F.Z., L.W. and X.C. performed co-registration between EM dataset and micro-CT dataset. B.H., L.W., H.Z., and J.G. annotated the ground truth dataset. J.L., B.H., J.Z.L. and Q.X. developed the dense reconstruction method. X.Z. provided the parallel computing environment. J.L., H.Z. and Y.Z. performed the large-scale segmentation and reconstruction of EM data. H.H., L.S., J.B.Y. and D.Z. developed and managed the annotation and proofreading software. L.M., Y.C., M.L. and Z.W. designed and developed the visualization pipeline, and generated the sub-figures and videos. J.L., J.Y., H.Z., Y.C., L.M., H.H. and H.C. designed the figure layout and finalized the figures. J.L. and J.Y. performed the data analysis under the supervision of H.H. and H.C.. Y.X. provided valuable suggestions on various aspects of the project. J.L., J.Y., L.S., X.C., L.M., H.H. and H.C. wrote the paper and all authors participated in the discussion, data interpretation and conception during the project.

## Materials and Methods

### Animals

All animal experiments were conducted in accordance with the guidelines of the Animal Care and Use Committee of Peking University accredited by AAALAC International and the procedures were approved by the Animal Care Committee of Peking University (Approval ID: IMM-ChengHP-14). Wild type C57BL/6J mice were purchased from Vital River Laboratory and maintained at a controlled environment (temperature: 22 ± 2℃, humidity: 60 ± 5%) on a 12-hour light/dark cycle, with water and food available *ad libitum*.

### Preparation of 1-mm^3^ SCN tissue block

Before the experiment, 6-week-old male mice weighing between 20 and 25 g were housed in individual cages with running wheels for 2 weeks, with 250-lux illumination during the light. The time of lights-on and lights-off were defined as ZT 0 and ZT12, corresponding to 07:00 and 19:00, respectively. The locomotor activities of the mice were recorded and analyzed by ClockLab (Actimetrics). Mice exhibiting robust behavior rhythms were selected for the subsequent preparation of SCN slice.

The mice were removed from their home cage at ZT11.5 and anesthetized with 1.25% tribromoethanol (20 μL/g, i.p.; EasyCheck). The perfusion procedure was performed as previously detailed(*1, 2*) and was conducted within 30 min before the time of lights-off. Briefly, the mice were first perfused with 0.9% saline, followed by 2.5% glutaric dialdehyde (GA, Sigma) in a 0.1 M sodium cacodylate trihydrate buffer (pH 7.4, Biolink). Subsequently, the mice were decapitated, and the brains were removed to a pre-fixative solution consisting of 4% paraformaldehyde (PFA, Sigma) and 2.5% GA in a 0.1 M sodium cacodylate trihydrate buffer at 4℃ over 12 hours. Note that the optic nerves should be carefully cut to prevent any undue stretching of the SCN. On the second day, the brain was transferred to ice-cold 0.1 M phosphate buffered saline (PBS, Sigma), and then a 1-mm-thick coronal slice containing the entire SCN was prepared using a vibratome (VT 1200S, Leica). Following this, a tissue block of the SCN (1 mm^3^) was meticulously separated using a pair of scalpels under a stereo microscope (Stemi 508, Zeiss).

### EM sample preparation

The preparation of EM samples followed the *en-bloc* staining procedure(*1*). First, the SCN tissue block was transferred to a glass tube fulfilled with 0.15 M sodium cacodylate trihydrate and washed 3 times (10 min each) at room temperature (RT, 22–25℃). Next, the tissue block was post-fixed in 2% OsO_4_ (buffered in 0.15 M sodium cacodylate trihydrate) (Ted Pella) for 90 min, followed by staining in 2.5% potassium ferrocyanide (Sigma) dissolved in 0.15 M sodium cacodylate trihydrate for another 90 min. After thorough washing with ddH_2_O (3 times, 20 min each), the tissue block was incubated sequentially with filtered thiocarbohydrazide (Sigma) at 40℃ for 45 min, 2% OsO_4_ (buffered in ddH_2_O) at RT for 90 min and 1% uranyl acetate (Merck) aqueous solution at 4℃ for 12 hours.

Subsequently, the tissue block was washed with ddH_2_O (3 times, 20 min each) was dehydrated through a graded ethanol series (50%, 70%, 80%, 90%, 100%, 10 min each), followed by an ethanol/acetone series (1:1, 0:1, 0:1, 0:1, 10 min each). The tissue block was then embedded through an acetone/epon-812 resin (SPI) series (1:1, 1:2, 1:3, 0:1, 3–4 hours each). Finally, the fully infiltrated tissue block was placed in an embedding mold containing pure resin at 37℃ for 24 hours, 45℃ for 24 hours and 60℃ for 24–48 hours, successively. Note that all steps were performed within glass tubes, which were replaced at each step to prevent the accumulation of solution residue.

Following the preparation of EM samples, micro-CT imaging (Xradia 515 Versa, Zeiss) was conducted on the SCN tissue block, capturing coronal, sagittal, and horizontal perspectives with a resolution of 1.2 μm/voxel to fully depict its internal structure. EM samples selected for subsequent ultra-thin serial sectioning were assessed against specific criteria to ensure the preservation of both morphological integrity and continuity of the SCN.

### Sample sectioning

The EM sample was sectioned and collected using an RMC Powertome ultramicrotome, in conjunction with an ATUMtome automated collection system (RMC)(*2*). Initially, the specimen surface was refined. The ATUMtome was then used to gather sections at the liquid level of the ultramicrotome diamond knives, transferring them onto Kapton sample-bearing rolls via a conveyor mechanism. The section thickness was set to 40 nm, and the cutting speed was set to 0.8 mm/s. During the cutting process, the knife was manually shifted or replaced when sections became fragmented.

For the bookbinding process of the slices, the rolls were cut to the required length and adhered to 10 cm diameter silicon wafers using double-sided carbon conductive tape. The assembly was then baked in an oven at 60℃ for over 30 min to remove excess moisture from the sections. Subsequently, the silicon wafers containing the slices were placed in a high vacuum coater for carbon spraying.

### Wafer fabrication and mapping

The adhered wafers were first subjected to carbon coating with a film thickness of 10 nm (ACE 600, Leica) to enhance the electrical conductivity of the sample surface and minimize charge accumulation. Next, custom-developed wafer pre-irradiation equipment was used to irradiate the entire wafer, improving sample stability and enhancing backscattered electron signal contrast(*3*). The irradiation was performed at a current of 9 mA and a voltage of 2000 V for a continuous duration of 10 s. The processed wafers were then placed under an optical microscope (Clippers, FBT) for the acquisition of navigation images, by line-scan rapid imaging with a resolution of 7.6 μm. We manually annotated the ROIs in each section, which were then used by a software to enable the automated acquisition workflow of an electron microscope.

### ssEM image acquisition

High-throughput image acquisition was conducted simultaneously across three scanning electron microscopes (Navigator NeuroSEM100, FBT) using custom-designed software to maximize imaging efficiency. The pixel resolution was set at 5 nm, with a dwell time of 43.5 ns and an acceleration voltage of 2 kV. Autofocus and astigmatic correction were applied for 80 s per section.

To expedite the acquisition rate and increase the probability of successful imaging, a methodical region-specific focusing and imaging strategy was employed. Firstly, the imaging ROI of each section was divided into multiple tiles (12k × 12k pixels) with approximately 10% overlaps. These tiles were then grouped into regions of 25 tiles arranged in a 5 × 5 grid. Regions with fewer than 25 tiles at the section edge were also considered as individual standard regions. These regions were numbered starting from ‘region1’ and follows a serpentine pattern based on their positions.

Prior to scanning each section, a low-resolution focus was conducted to ensure the algorithm’s efficacy in identifying the optimal focal range at the center of the section, set at a resolution of 50 nm with a dwell time of 80 ns. Subsequently, the stage was precisely oriented to the center of the first region, where high-resolution focusing and astigmatic correction were performed at a resolution of 5 nm with a dwell time of 80 ns. Following focusing, imaging proceeded in a snake-like trajectory for each tile. Upon completion of a region, the stage moved to the next region’s position, continuing the high-resolution focus and scanning process. The ideal stage movement and data transfer time was 2.5 s per tile. This cyclical pattern continued until all regions of the current section were captured, with transitions between regions following a serpentine navigation pattern. This process was repeated for subsequent sections until the entire wafer was imaged.

In parallel, a defocus detection module was used to ensure the image quality of tiles. The defocus detection module used the statistical gradient feature to measure the image blurring. We first compared the gradient value of the overlapping regions between adjacent tiles, and then employed maximum spanning tree to evaluate the relative blurriness. By combining automated detection with manual inspection, low-quality images were identified and reimaged.

### Section loss

Due to the ultra-thin nature of the sections and the characteristics of resin materials, issues such as contamination, cracks, folds, and even section loss are unavoidable during sectioning, collection, knife changes, and imaging. To mitigate the impact of these issues on subsequent processes, a comprehensive manual inspection was conducted on the entire SCN dataset.

The knife-changing or knife-shifting process resulted in the loss of 38 sections, which were not imaged due to the absence of imaging ROIs. An additional 22 sections with partial ROIs were imaged and included in the SCN dataset. Furthermore, 38 sections exhibited significant cracking or folding issues. To maximize the retention of useful information outside the damaged areas, these sections were repaired using the Expected Affine algorithm(*4*).

### Stitching and alignment

A total of 6,824 serial sections were selected to compose a 3D image stack. Each section consisted of ∼96 tiles (12 rows × 8 columns), each with a size of 12k × 12k pixels. First, the tiles of each section were stitched together. Then, a coarse-to-fine alignment pipeline was used to align the serial sections.

### Stitching

The serial sections were imaged as partially overlapping tiles (∼10% overlap) by moving the microscope stage. In practice, approximately 200**–**500 Scale Invariant Feature Transform (SIFT)(*5*) key point pairs were extracted from the overlapping regions between adjacent tiles. These key point pairs were used to calculate the relative positional relationship between two adjacent tiles. A least squares method was then applied to globally optimize the position of each tile, laying a crucial foundation for the subsequent seamless stitching processes.

### Alignment

We present our registration pipeline for stitched serial sections. The original sequence was divided into 228 “stacks” with a one-section overlap. A “stack” refers to a group of 31 consecutive sections from the original sequence, serving as the primary unit for subsequent coarse-to-fine alignment. To ensure structural continuity across all stacks, the first section of each stack was sampled, resulting in a sequence of 228 reference sections. Affine transformations were applied to correct significant deformations in the down-sampled reference section sequence. Each pair of adjacent reference sections was then positioned at the beginning and end of each stack, with the remaining sections within each stack adjusted through the coarse and fine registration process. The reference sections in every stack remain consistent throughout the overall registration.

During the coarse registration process, reference sections were down-sampled by a factor of 32. Pairwise coarse registration was performed by extracting correspondences from two adjacent sections and applying affine transformation to deform sections, gradually registering the entire stack. The extraction of the feature points and computation of raw correspondence between adjacent sections were performed using CudaSift(*6*), which implements the SIFT algorithm(*5*) on a GPU for high throughput and minimal latency. Raw correspondences were then filtered using the RANSAC algorithm(*7*) on a CPU to enhance reliability. After above calculations, approximately 500 correspondence pairs remained for each section pair. These correspondences were used to compute the affine transformation between two adjacent down-sampled sections, and the original sections were deformed accordingly.

For fine-scale registration, we optimized the elastic method(*8*) on images down-sampled by a factor of 16 from the original sections. The elastic algorithm searches for dense correspondences by calculating the Normalized Cross-Correlation coefficients between patches on adjacent sections across the entire down-sampled stack. The algorithm then optimized the elastic deformation mesh based on dense correspondences. The original, unsampled sections were subsequently deformed using thin-plate spline(*9*) based on the up-sampled optimized meshes. Finally, the deformed sections in stacks were ultimately exported as overlapping tiles for subsequent reconstruction.

### Co-registration between ssEM and micro-CT data and identification of SCN boundary

The capillaries were used as the bridge for co-registration between ssEM and micro-CT data. Initially, we manually annotated the fragments of large capillaries in micro-CT data using the TrakEM2 plugin in Fiji(*10*), and automatically detected all the capillaries in ssEM data using a 3D Convolutional Neural Network (CNN) (see section of capillary detection). Nine distinct trinomial or quadrinomial points of capillaries were manually selected from each dataset as alignment markers. An affine transformation matrix was computed based on these points to achieve the initial alignment. The alignment was further refined using the Iterative Close Point algorithm(*11*), with a maximum iteration count of 100k to ensure precise alignment. This iterative process ultimately yielded a deterministic affine matrix.

Micro-CT data, with its isotropic resolution and wide field of view, enables a more accurate delineation of the SCN contour. Thus, we first delineated the SCN contour in the micro-CT data using the Vaa3D(*12*), and then transformed it into the ssEM space by applying the computed affine matrix.

### Valid tissue and defect detection

Image defects and valid tissue masks were precomputed for subsequent detection(*14*), (*15*). Based on the grayscale features of the SCN EM images, valid tissue masks were identified using a simple thresholding method. We combined manual labeling with CNN-based methods for defect detection. Six sub-volumes (each size of 3200 × 3200 × 5 pixels) were annotated as ground truth for defect detection. A 2D UNet was trained to classify every voxel as either third ventricle (3V), capillary or membrane-damaged region at a voxel size of 40 × 40 × 40 nm^3^. The input patch size was 385 × 385 pixels. The network was trained for 500k steps using a weighted cross-entropy loss and a Stochastic Gradient Descent (SGD) optimizer with a learning rate warmup strategy (learning rate of 0.04, linear warmup of 2k steps, cosine decay). Following this detection, annotators further labeled regions affected by folds, cracks, and artifacts resulting from the diamond knife replacing process. All the aforementioned masks together constitute the defect masks.

### Local realignment

Local misalignments are inevitable in large-scale ssEM stacks, negatively impacting segmentation accuracy. Inspired by the Flood-Filling Network segmentation pipeline(*13*), we proposed realigning the local sub-volumes before CNN inference and segmentation. For each sub-volume, a 4 × 4 affine transform matrix was estimated. The sub-volume was then padded and transformed according to this matrix. After CNN inference, agglomeration or connected components labeling algorithms, the result was de-aligned and cropped back to its original shape (fig. S6A). To improve efficiency, affine transform matrices for all sub-volumes were precomputed and saved.

### Network architecture and Data augmentation

The networks used for segmentation tasks in this paper are 3D Residual U-Net^52^. This architecture includes an analysis path with four down-sampling levels and a synthesis path with four up-sampling layers. It employs summation-based skip-connections to integrate higher-level contextual information with lower-level localization details. Each level of the network consists of three convolutions (with SAME padding), followed by batch normalization and a Rectified Linear Unit (ReLU) activation function, all connected through residual skip connection.

All training processes were conducted on three NVIDIA Tesla V100 GPUs. During training, training samples were randomly augmented by various transforms, including misalignment (up to 16 pixels), missing sections (up to 2 slices), out-of-focus section (up to 2 slices), random rotations, flips and grayscale transformations^52^.

### Block-based strategy for large-scale network inference

A block-based strategy was designed for parallel network inference on the full SCN dataset. The original raw images were divided into sub-volumes with overlaps, which were distributed across 14 nodes of a spark cluster, each equipped with three NVIDIA GeForce RTX 3090 GPUs. Each sub-volume was first padded and realigned, then cropped into small patches with overlaps. After CNN inference, the overlapping probability maps were stitched using a Gaussian bump function^52^. The final predictions for each sub-volume were then de-aligned, cropped, and saved.

The block and overlap sizes were set to 5256 × 5256 × 108 pixels and 128 × 128 × 4 pixels at a resolution of 10 × 10 × 40 nm^3^ for affinity map, synapse, mitochondria and DCV/SSV detection task; 1506 × 1506 × 108 pixels and 128 × 128 × 4 pixels at a resolution of 40 × 40 × 40 nm^3^ for nucleus detection task; 881 × 881 × 108 pixels and 128 × 128 × 4 pixels at a resolution of 80 × 80 × 40 nm^3^ for capillary, axon fascicle and soma detection task.

### Block-based connected components labeling pipeline

Similar to the block-based strategy for network inference, we also designed a block-based connected components labeling (CCL) pipeline for parallel 3D labeling. The entire dataset was divided into sub-volumes, which were distributed across 14 nodes of a spark cluster, each equipped with 20 CPUs. Each sub-volume was first padded and realigned, followed by the application of the CCL algorithm for 3D instances. The resulting 3D instance map of each sub-volume was then de-aligned, cropped, and saved. Once all sub-volumes were processed, each pair of adjacent sub-volumes were merged based on the shared face, forming the final composite 3D instance map.

The block size was set to 5000 × 5000 × 100 pixels at a resolution of 10 × 10 × 40 nm^3^ for synapse and mitochondria CCL pipeline; 1250 × 1250 × 100 pixels at a resolution of 40 × 40 × 40 nm^3^ for nucleus CCL pipeline; 625 × 625 × 100 pixels at a resolution of 80 × 80 × 40 nm^3^ for capillary, axon fascicle and soma CCL pipeline.

### Dense reconstruction

The SCN dataset was densely segmented using a boundary detection-based method(*16, 17*). A 3D CNN was used to predict three nearest-neighbor affinity maps, which were then post-processed with the distance transform watershed (DTW) algorithm(*16*) to generate super-pixels for each slice. Agglomeration based on region adjacency graph (RAG) was used to merge over-segmented neuronal fragments (fig. S6).

### Affinity map

To train an affinity predictor, experienced human experts manually annotated 5 sub-volumes containing 145,162,600 voxels at a resolution of 10 × 10 × 40 nm^3^. A 3D Residual U-Net was trained to predict three nearest-neighbor affinity maps on the entire SCN dataset. The input patch size was 256 × 256 × 8 pixels. The network was trained using binary cross-entropy loss and an SGD optimizer for 500k iterations. The learning rate was initialized at 0.01 and decayed by a factor of 0.1 at iteration steps 30k and 35k.

### Intra-chunk agglomeration

To construct a signed graph from the predicted affinity maps, we first performed DTW on each 2D slice (boundary map averaged across the three channels of affinity maps) and relabeled the results to guarantee each 2D super-pixel had a unique ID. Then, a 3D RAG was extracted and two sets of edges (intra-slice edges and inter-slice edges) could be determined. 191-dimensional edge features were extracted from statistics of raw images and boundary maps of the corresponding boundary and region(*16*). Using these edge features, we trained two random forest classifiers to independently learn scores for two types of edges. A score of 1 represents a repulsive edge score, while a score of 0 indicates an attractive edge. The predicted scores were then transformed into costs using the following formula:

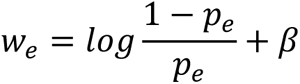

where *p_e_* denotes the edge score, and *β* is the boundary bias, which adjusts the degree of over-segmentation. We set *β* to 0.6 in this dataset. We then employed a greedy agglomerative clustering algorithm with the “mean” linkage criteria(*18*) to solve the graph partitioning problem. The agglomeration was performed on sub-volumes of 2700 × 2700 × 52 pixels with overlaps of 100 × 100 × 1 pixels. Before agglomeration, each sub-volume was padded and realigned. After agglomeration, the final segmentation result was de-aligned, cropped, and saved. The overlapping area was saved separately for further processing.

### Inter-chunk matching

To merge segments between adjacent chunks, we identified all matching segment pairs using a custom overlap-based method (fig. S6, B and C) designed to prevent merging in regions with image defects. These pairs were then merged to generate the final global segmentations. For each chunk pair (denoted as *A* and *B*), the valid region masks and defect masks were prepared for the following step. Every two overlapping segments (denoted as *a* and *b*) were then determined whether to merge or not according to the following rules:

1. If the number of invalid slices exceeds 6 in either *A* or *B*, all segment pairs between *A* and *B* are excluded from merging.
2. If the proportion of the 3V region exceeds 30% or the membrane-damaged region exceeds 50% in either *A* or *B*, no segment pairs between *A* and *B* are allowed to merge.
3. If the Intersection over Union of *a* and *b* in the shared overlapping area exceeds 70% and the number of overlapping voxels exceeds 40,000, the pair *a*-*b* is considered for merging if it satisfies condition 5.
4. If either overlap ratio exceeds 90% or both overlap ratios exceeds 80%, the pair *a*-*b* is considered for merging if it satisfies condition 5.
5. If either segment *a* or *b* contains defects, the corresponding overlapping ratio must reach 90%; If both segments contain defects, the overlapping ratio for both must reach 98%.

### Capillary, fascicle and nucleus detection

Capillaries and axon fascicles were automatically segmented by applying a separately trained 3D Residual U-Net to the full dataset at a down-sampled resolution of 80 × 80 × 40 nm^3^. Five sub-volumes were manually annotated as the ground truth for each target, covering 62,202,275 voxels for capillaries and 597,297,500 voxels for axon fascicles. The input batch size was set to 256 × 256 × 8 pixels. Both networks were trained using binary cross-entropy loss and SGD optimizer, with a learning rate strategy incorporating a linear warmup of 2k steps and cosine decay. The initial learning rates were set to 0.02 and 0.01, with total training steps of 300k and 100k, respectively. After inference, a block-based CCL method was then applied to the thresholded map (0.78 for capillary and 0.4 for axon fascicle), resulting in the final detections of capillary and fascicle. Axon fascicles of length greater than 200 μm were used for analysis.

Nuclei were detected at a resolution of 40 × 40 × 40 nm^3^ (fig. S7A). Experienced annotators manually labeled all nuclei within eight sub-volumes of 687 × 687 × 55 pixels as ground truth. A 3D Residual U-Net was then trained to predict the probability of each voxel belonging to the foreground. The input batch size was set to 256 × 256 × 8 pixels. The network was trained using binary cross-entropy loss and SGD optimizer, with the learning rate initialized at 0.01 and decayed by 0.1 at iteration step 30k, 50k and 70k. The total training iteration was set to 500k. Following thresholding of the full-size probability maps at a threshold of 0.7, a block-based CCL algorithm was employed to obtain 3D instances of nuclei.

### Soma detection, postprocessing and graph construction

Soma segmentation and postprocessing were performed at a down-sampled resolution of 80 × 80 × 40 nm^3^ (fig. S7A). We annotated five sub-volumes, covering regions containing somata as well as neurons, glia, and capillaries, totaling 335,554,900 voxels, which were used for training and testing. Direct membrane contact between somata is common in the SCN, so we employed hybrid representation learning(*19*) to predict binary foreground, contour, and distance map for each soma instance using a residual 3D Residual U-Net(*17*) with three-channel outputs, aiming to prevent potential merge errors. The soma detection model was trained using an SGD optimizer with a momentum of 0.9 and a weight decay of 0.001 for 100k iterations. Linear warmup and cosine annealing were used in the learning rate scheduler, starting with an initial value of 0.02. The input size was set to 256 × 256 × 32 pixels. Evaluation against the test set yielded a precision of 92% and a recall of 92%. During the inference phase, the consensus between the representations of foreground and boundary was subjected to thresholding at a threshold of 0.78, which then served as input for the block-based CCL pipeline.

Given the nucleus detection, we first leveraged the correspondence between the nuclei and somata segmentations to identify soma merge errors. We then performed local realignment on the merged soma regions. Using the nuclei as initial seed points, we applied the marker-controlled watershed transform to the segmented foreground, effectively separating the somata. The separated somata were then de-aligned and saved.

The construction of the soma-soma ephaptic interaction network was performed at a resolution of 160 × 160 × 160 nm^3^. We iterated through all the somata, treating each soma as a node in this graph. If contact was detected at the boundary between two somata, an edge was established between the corresponding nodes. Specifically, we traversed the boundary pixels of each soma; if another soma (*B*) was detected within the horizontal four-connected components of the boundary pixels of soma (*A*), the current boundary pixel was considered to be on the contact surface between soma *A* and *B*. The contact area between the somata was quantified and assigned as the weight of the edge. The undirected weighted graph was then decomposed into multiple EIUs by solving the minimum cut problem^41^, with a bias parameter set to 0.85.

### Graph property of the ephaptic interaction network

To characterize properties of the soma-soma ephaptic interaction graph, we used two types of random graphs as the null model. The ER model(*20*) is a type of random graph in which any pair of nodes is connected with equal probability. The CFG preserves predetermined degree sequences by applying a double-edge swap algorithm(*21*) to the given graph. Rich club is a kind of graph property(*22*), the high-degree nodes tend to form tightly interconnected clubs. The rich club coefficient measures the presence of rich club, is defined as a function of node degree:

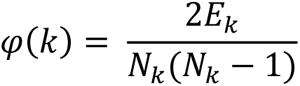

where *E_k_* is the number of edges when nodes with degree greater than *k* are removed. *N_k_* is the number of nodes with degree larger than *k*. To compare the tendency relative to CFG, the relative rich club coefficient was normalized by randomly sampling 1,000 CFGs. We used both the ER and CFG models to study the frequency of three-node motifs in the soma-soma ephaptic interaction graph. For each actual EIU, the corresponding ER and CFG models were repeatedly sampled 100 times, and the number of motifs was computed and compared with the actually observed value.

### Classification of cell types

We trained a random forest classifier to categorize cell types within the SCN. Each soma was paired with the nucleus that had the highest overlap. From both the nucleus and soma, we extracted geometric and texture features, forming a 46-dimensional feature vector. We manually labeled 328 endothelial cells, 457 glia and 212 neurons as ground truth based on their distinct 3D morphologies and somatic appearances in EM images. The dataset was then divided into a training set (80%) and a test set (20%). The trained classifier achieved an average accuracy of ∼89% on the test set and was subsequently applied to all detected somata for cell type classification. However, 203 objects remained unassigned due to segmentation errors.

### Automated synapse detection and assignment

We trained a 3D Residual U-Net to identify binary masks of synapses at a resolution of 10 × 10 × 40 nm^3^ (fig. S7B). We annotated 877 chemical synapse instances within nine sub-volumes, totaling 4,502 μm^3^, as ground truth. Six sub-volumes were used for training and the rest for testing. SGD was applied to optimize the synapse detection network. A learning rate strategy with linear warmup (2k steps) and cosine decay (starting from 0.01) was used. During training, we used weighted binary cross-entropy loss and dice loss, with foreground voxels weighted according to the ratio of positive to negative voxels. We trained the network for 500k iterations with input batches of 256 × 256 × 8 pixels. Evaluation against the test set yielded a precision of 91% and a recall of 85%. The resulting probability maps were binarized at a threshold of 0.5, and a block-based CCL algorithm was applied to obtain 3D synapse instances. Synapses smaller than 500 voxels were filtered out as false positives.

To identify the pre-synaptic and post-synaptic neurons for each synapse, we employed another 3D CNN to predict the local pre/post-synaptic neuronal masks(*15*) at a resolution of 10 × 10 × 40 nm^3^ (fig. S7B). The ground truth dataset contains 222 synapses. Training samples were centered on each synapse and padded with 0 if they were near the boundary. The input size of the network is 80 × 80 × 18 pixels. Binary cross-entropy loss was used for both output channels. The model was trained using the SGD optimizer with a learning rate of 0.01 for 100k steps. The trained network was then applied to all detected synapses, and the synaptic partners were identified by assigning neuronal segment IDs with which the predicted masks (using a threshold value of 0.9) had the most overlap.

### Automated detection of mitochondria, DCVs and SSVs

We detected the mitochondria using a trained 3D Residual U-Net at the resolution of 10 × 10 × 40 nm^3^ (fig. S7B). 986 mitochondria across 3 sub-volumes, covering 933 μm^3^, were manually labeled as the ground truth. The output layer was designed for two tasks: predicting the mitochondrial inner mask and the mitochondrial membrane. Binary cross-entropy loss was used for both tasks of the mitochondria detection network, with an additional dice loss applied to the mitochondrial membrane task. The network was trained for 500k steps using the SGD optimizer. The learning rate was initialized as 0.01, with 2k linear warmup steps, and manually decayed at step 30k, 50k and 70k. The input size was set to 256 × 256 × 8 pixels. The final mitochondria probability map was generated by taking the intersection of the binarized inner map (thresholding at 0.65) and the binarized mitochondrial membrane map (thresholding at 0.5). A block-based CCL algorithm was ultimately used to reconstruct the mitochondria instances.

A three-class semantic segmentation model was used to classify each voxel as either DCV, SSV, or background (fig. S7B). Due to the limited resolution, DCVs and SSVs were annotated as either “clustered” or “scattered” within 6 sub-volumes, rather than individual instances. The ground truth encompasses various structures, totaling 236,937,520 voxels at a resolution of 10 × 10 × 40 nm^3^. The input batch size was set to 256 × 256 × 8 pixels. The vesicle detection network was trained for 500k iterations using cross-entropy loss and the SGD optimizer. A learning rate strategy with linear warmup (2k steps) and cosine decay (starting from 0.01) was applied. The trained network yielded a precision of 91% and a recall of 84% for DCVs, and a precision of 91% and a recall of 95% for SSVs. During the inference phase, each voxel was assigned to the class with the highest predicted probability.

### Spatial distribution maps of multi-scale reconstructions

To visualize spatial fluctuations of reconstructed attributes across the SCN, we computed the local parameters by evenly gridding the SCN space with gird sizes of 10 μm for the density and mean size of synapses and mitochondria; 10 μm for the volume fraction of DCVs and SSVs; 20 μm for the density and mean size of somata; and 30 μm for the density of branch points and mean radius of capillaries. For linear correlation analysis, the local densities of all attributes are computed within gird sizes of 10 μm to ensure consistency.

### Manual skeleton labeling and proofreading

ACE (Advanced Connectome Engine) is a specialized software solution tailored for the reconstruction of large-scale connectomic data. It integrates massive data management capabilities, intelligent computing algorithms, and collaborative tools. Utilizing its engineering-driven process management, ACE guarantees efficient and accurate reconstruction of connectomes.

A multi-resolution SCNEM project dataset, based on a pyramid model, was generated using the ACE for manual neuron labeling and proofreading. We initially annotated the skeletons of 194 neurons, selecting their somata from approximately three z-planes. Skeleton points were meticulously traced and labeled on every z-plane, starting from the soma. Each skeleton was annotated by one individual and then reviewed by another to ensure accuracy and integrity. The skeleton annotation process required a total of 2,749 work hours. For proofreading the segmentation of each skeleton, the corresponding membrane contour were computed and imported into ACE according to the dense automatic segmentation results. We corrected errors in the contours by traversing each skeleton point, and the entire contour correction process took 3,698 work hours.

In addition to the skeleton-based proofreading, we randomly selected 197 neurons across the entire SCN for coarse proofreading. Dense reconstruction results of these neurons were imported into ACE. Annotators scanned the somata and dendrites of these neurons, correcting all the split and merge errors. Proofreading these neurons took 3,520 work hours.

### Skeletonization, compartment labeling and dendritic parameter

We used a distributed skeletonization method for the proofread neurons at a resolution of 40 × 40 × 40 nm^3^. The parameters of “scale” and “const” within the base TEASAR algorithm(*23*) were set to 4 and 10, respectively. To stitch the fragments within the skeleton of a single neuron, we connected the nearest nodes between subgraphs. Skeleton nodes with a radius greater than 2,000 were identified as soma and then aggregated into a single soma node based on their convex hull. The post-processed skeletons were imported into neuTube(*24*), a software tool for skeleton editing. Two annotators manually labeled the axonal and dendritic compartments for each neuron using a cross-validation approach.

The geodesic length was calculated as the total path length of the dendritic skeleton. For the axial dendritic domain, we projected the dendritic arbors along mediolateral, anteroposterior and ventrodorsal axes, and calculated the distance between the two extremes as its axial projection distances. The axial dendritic domain was determined by summing these three axial projection distances.

### Orientation and cross-section of axon fascicles

The automatically detected axon fascicles were transformed into skeletons using a distributed skeletonization method at a resolution of 40 × 40 × 40 nm^3^. The parameters of scale and const within the base TEASAR algorithm(*23*) were set to 40 and 300, respectively. Twigs shorter than 30 μm were pruned, and the skeletons were resampled into 10 μm segments. For each resampled skeleton point, the orientation was determined by calculating the direction vector of a line fitted through the three nearest neighboring points. The cross-section was defined as the intersection of the axon fascicle with the vertical plane.

### Quantification and statistical analysis

Non-parametric tests were used for all statistical analyses. The number of independent experiments (n) and the relevant statistical parameters for each experiment (such as mean ± s.e.m.) are described in the figure legends. Data fitting and calculation of the fitting coefficient (R^2^) were performed with Prism (GraphPad) and Origin 2021. The correlation coefficients *r* and *p* values were obtained from a two-tailed, non-parametric Pearson correlation analysis. In Boxplot, whiskers represent the minimum and maximum data points within 1.5 times the interquartile range from the first and third quartiles. Statistical significance was assessed with Prism, and the *p* value was calculated by two-tailed Mann–Whitney *U*-test. Significance levels are denoted as follows: ns: *p* > 0.05, \**p* < 0.05, \*\**p* < 0.01 and \*\*\**p* < 0.001, where applicable.

---

**Fig. S1.**
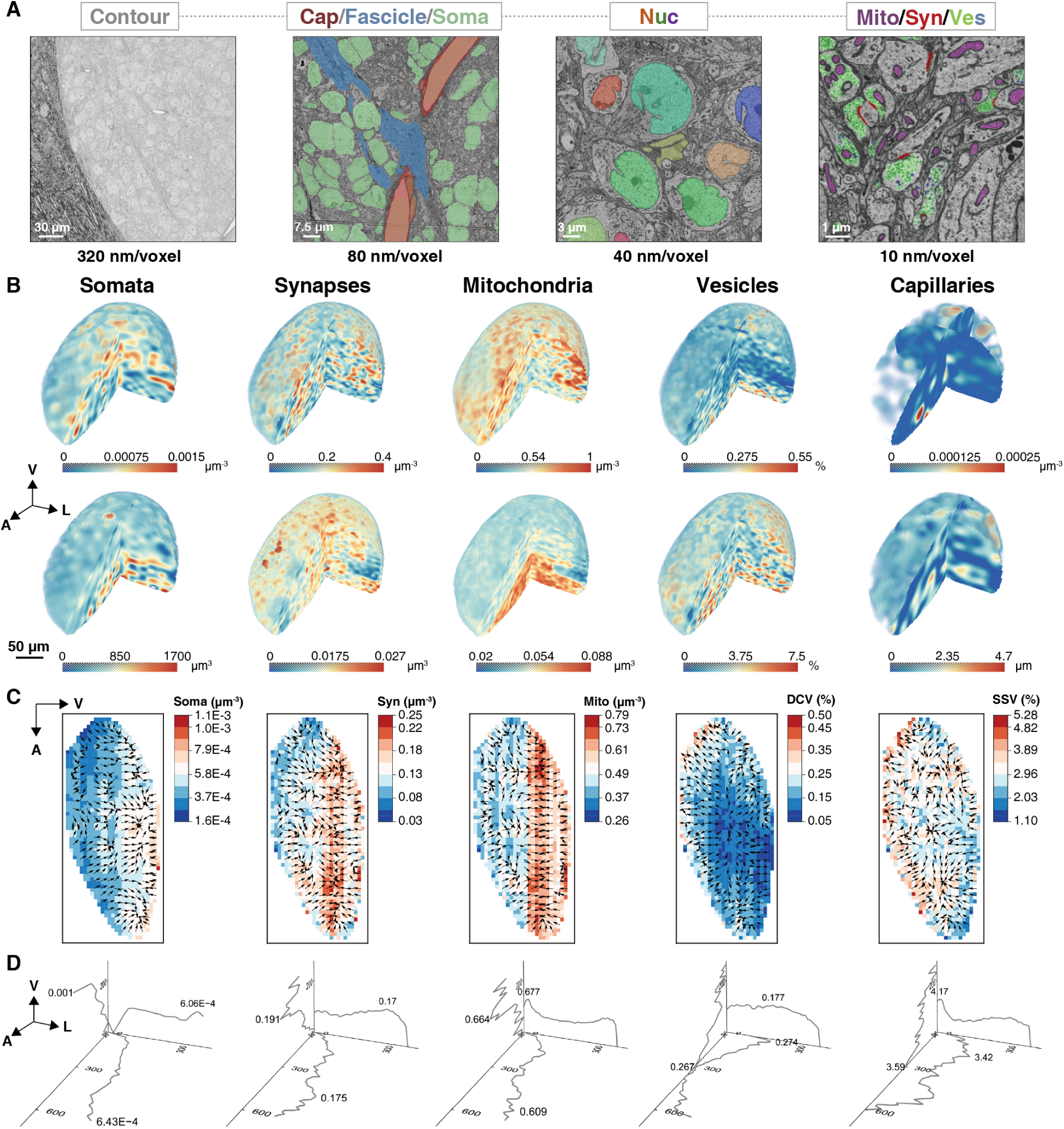
Multiscale and multimodal reconstructions of the SCNEM. (**A**) Multiscale reconstructions at different lateral resolutions in SCNEM dataset. Cap, Capillaries. Nuc: Nuclei. Mito: Mitochondria. Syn: Synapses. Ves: Vesicles. Scale bars, 30 μm, 7.5 μm, 3 μm and 1 μm. (**B**) 3D maps of substructures in the SCN. Somata, synapses, and mitochondria are shown by density (top, number per cubic micron) and mean size (bottom, mean volume per cube). Dense core vesicles (DCVs, top) and small synaptic vesicles (SSVs, bottom) are shown by volume fraction per cube. Capillaries are shown by branch point density (top, branch point number per cubic micron) and mean radius (bottom, mean radius per cube). Scale bar, 50 μm. (**C** and **D**) Averaged sectional density maps in the sagittal view (**C**) and axial density plots (**D**) of somata, synapses, mitochondria, DCVs and SSVs. Arrows in (**C**) represent the gradient directions. Cube sizes are as follows: somata: 20 × 20 × 20 μm^3^; synapses, mitochondria, DCVs and SSVs: 10 × 10 × 10 μm^3^; capillaries: 30 × 30 × 30 μm^3^. L, lateral; V, ventral; A, anterior.

**Fig. S2.**
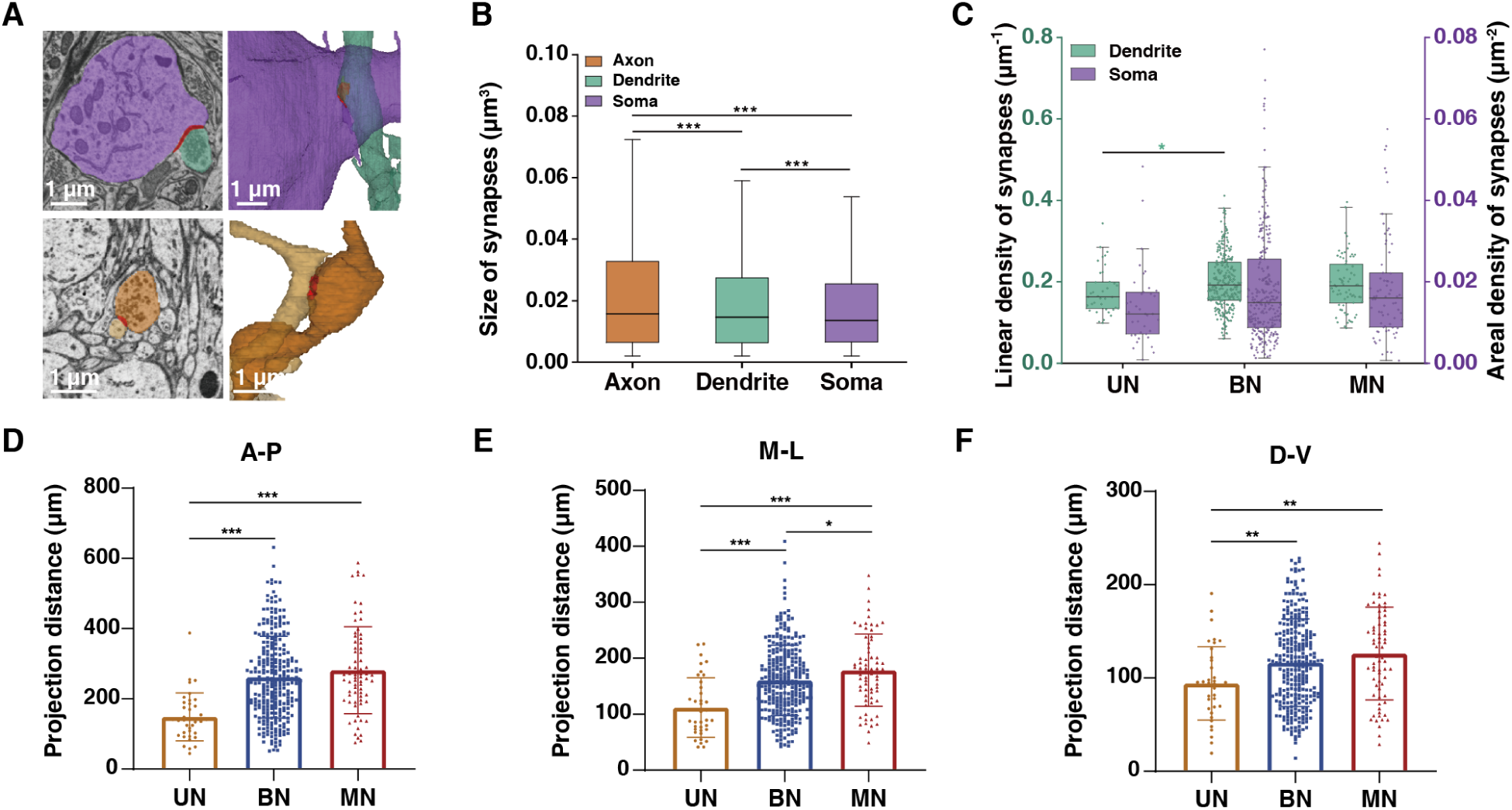
Characteristics of SCN neuronal attributes. (**A**) Representative EM images and 3D renderings of representative dendro-dendritic synapse (top) and axo-axonic synapse (bottom). Scale bars, 1 μm. (**B**) Size of synapses located on different neuronal compartments. Black lines denote the medians. n = 6,926, 6,276 and 144,368 in axon, dendrite and soma, receptively. (**C**) Linear density on dendrites and areal density on somata, computed as the number of synapses per 1 μm path length and per 1 μm^2^ surface area, respectively. n = 37 (UN), 284 (BN) and 70 (MN). (**D** to **F**) Dendritic projection distance along A-P axis (**D**), M-L axis (**E**) and D-V axis (**F**). A-P, anterior to posterior. M-L, medial to lateral. D-V, dorsal to ventral. n = 37 (UN), 284 (BN) and 70 (MN).

**Fig. S3.**
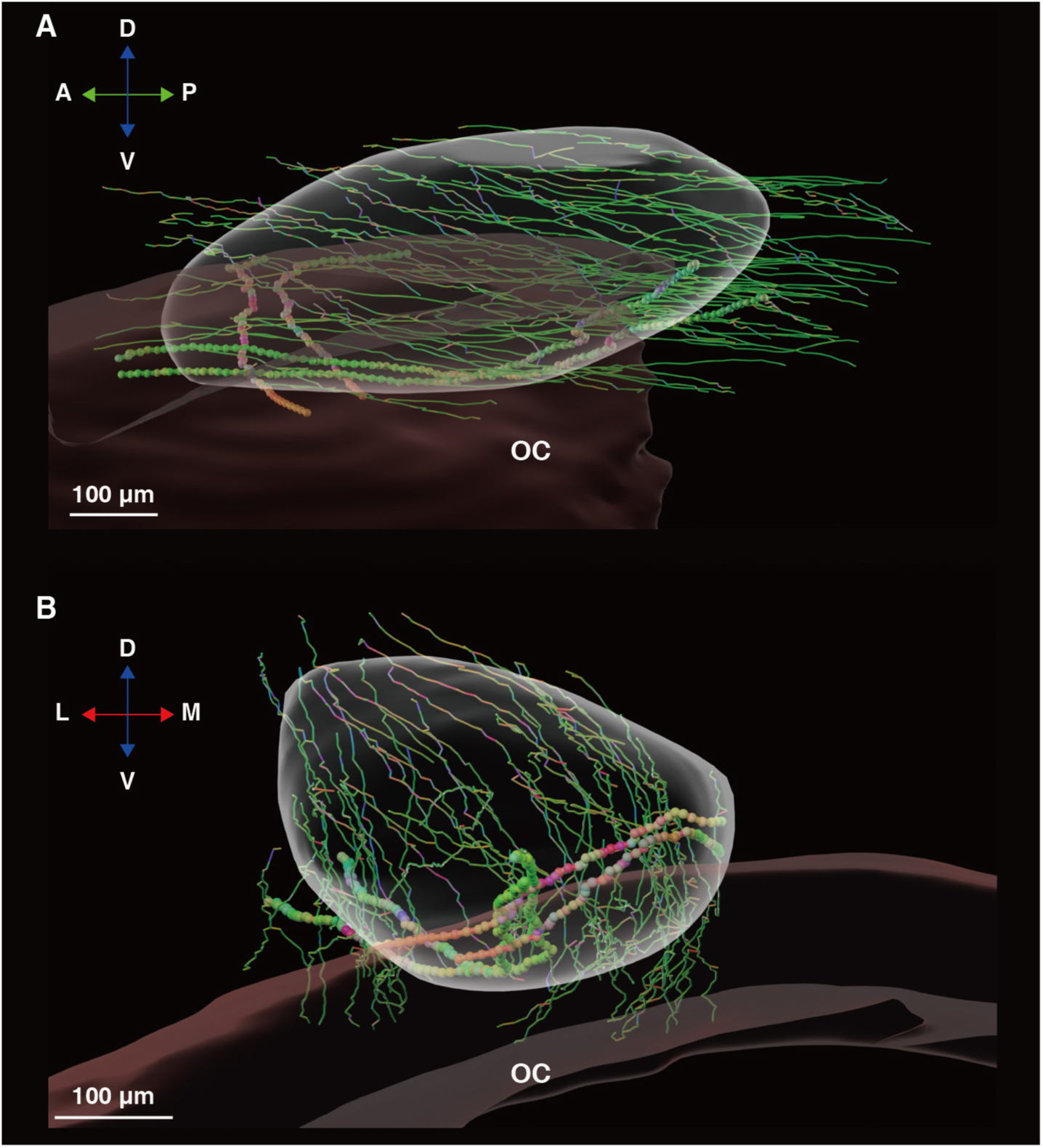
Trajectories of axon fascicles across the SCN and myelinated axons extending from the optic chiasm in sagittal (A) and coronal view (B). Anteroposterior, mediolateral, and ventrodorsal directions are colored in green, red, and blue channels, respectively. The radii of skeletons of axon fascicles and axons from optic chiasm are set to a ratio of 4:1. Scale bars, 100 μm. OC, optic chiasm; M, medial; L, lateral; P, posterior; A, anterior; D, dorsal; V, ventral.

**Fig. S4.**
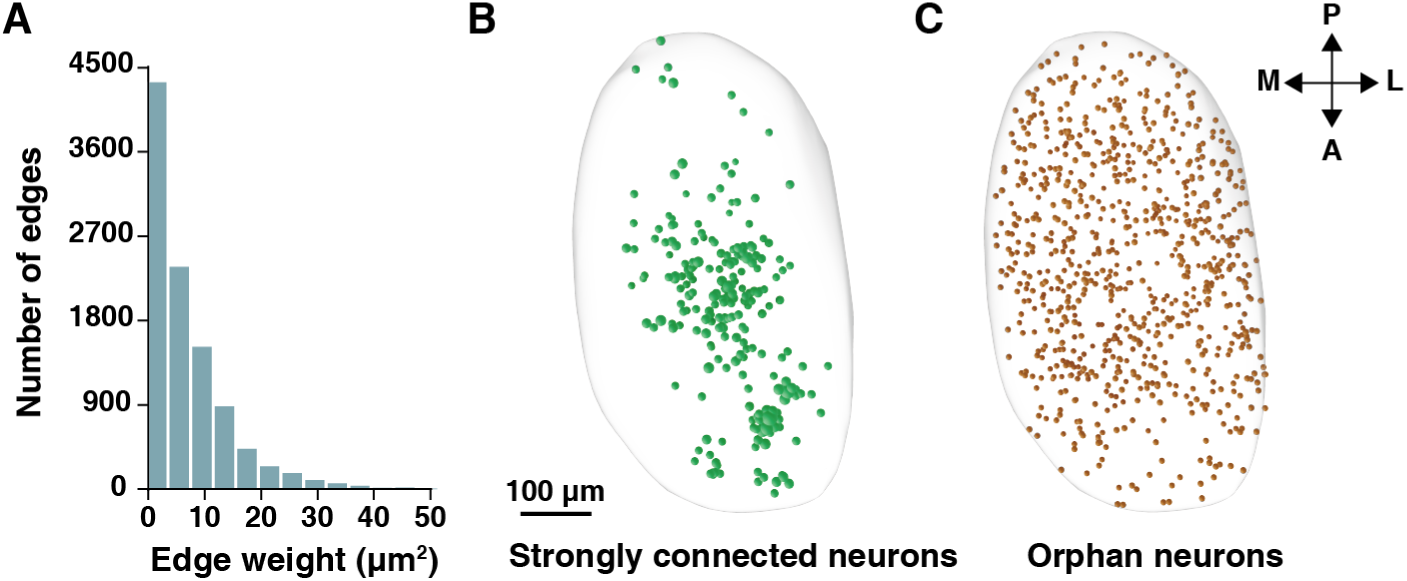
Properties of EIU network. (**A**) Distribution of edge weight, computed as the contact surface area between two adjacent somata. n = 10,169 edges. 6 observations with edge weight greater than 50 μm^2^ are omitted from the plot. (**B**) Spatial distribution of strongly connected neurons (green, with node degrees greater than five). (**C**) Spatial distribution of orphan neurons (orange, with node degrees equal to zero). Note that the size of node reflects the node degree. Scale bar, 100 μm. M, medial; L, lateral; P, posterior; A, anterior.

**Fig. S5.**
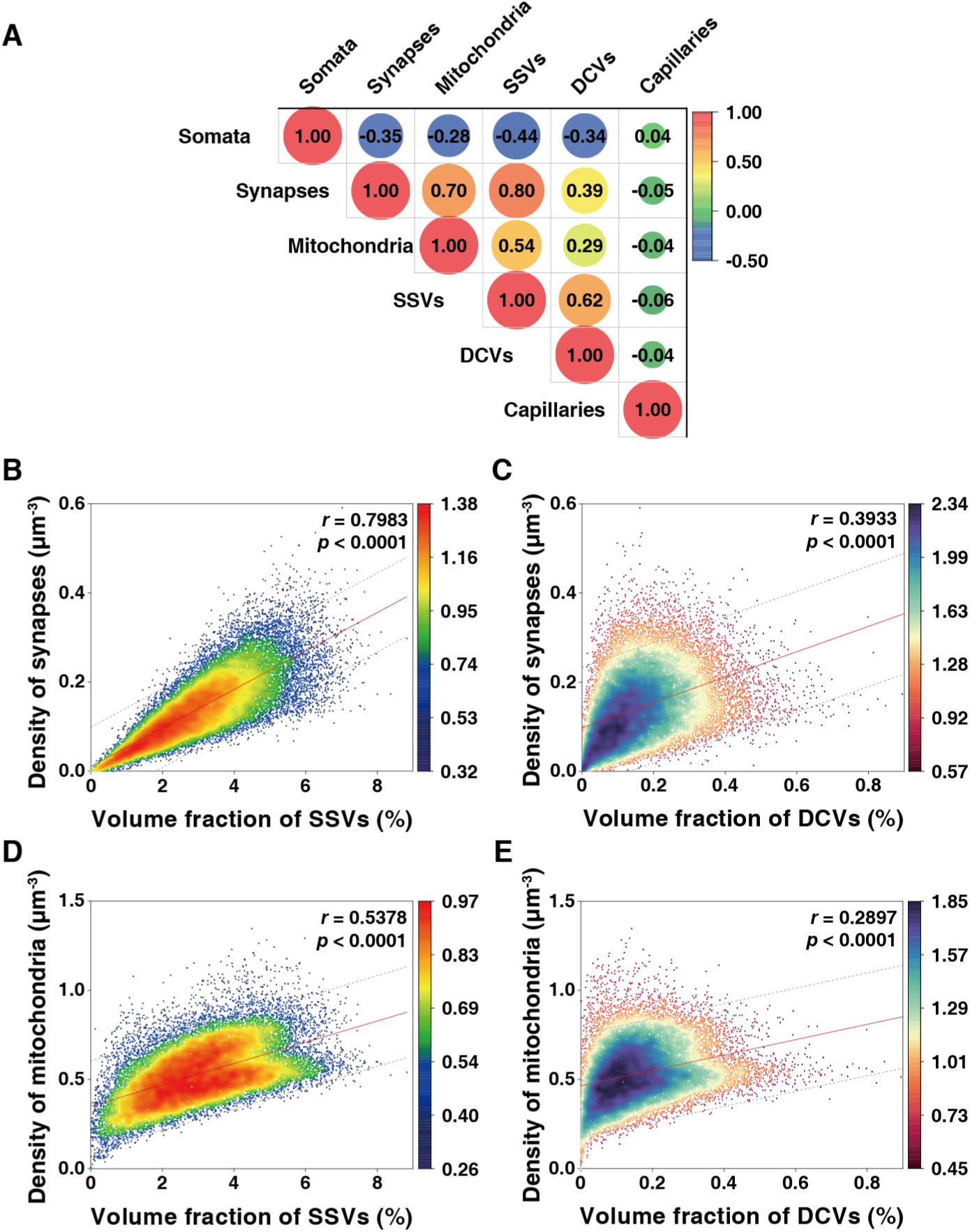
Cross-modality correlations. (**A**) Trans-modal correlation matrix analysis of spatial fluctuations of SCN parameters shown in **fig. S1, B**. Color code for Pearson’s correlation coefficients is shown on the right. (**B** to **E**) Linear correlation analysis between densities of synapses and SSVs (**B**) or DCVs (**C**), densities of mitochondria and SSVs (**D**) or DCVs (**E**). n = 30,083 points, each point represents the density computed within a cube of 10 × 10 × 10 μm^3^. Color codes of point density are shown on the right. In each panel, the red line represents the best-fitted linear function, and the grey dashed lines indicate the 95% prediction band. The correlation coefficients *r* and *p* values were obtained from a two-tailed, non-parametric Pearson correlation analysis.

**Fig. S6.**
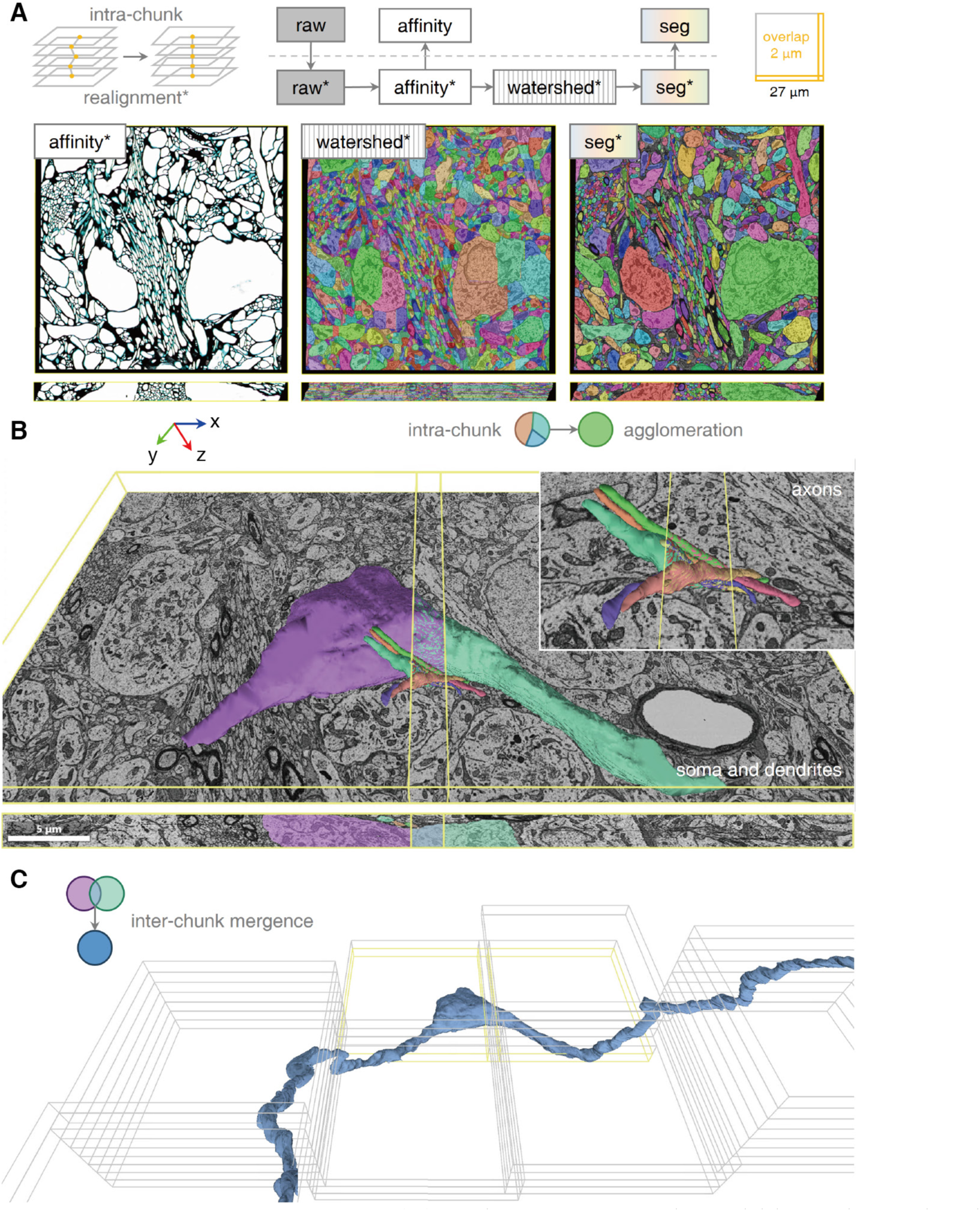
Pipeline for dense reconstruction. (**A**) Volume segmentation within each overlapping chunk (chunk size: 2,700 × 2,700 × 52 voxels, overlap size: 100 × 100 × 1 voxels). Before processing, each chunk (raw) was locally realigned (raw*) using affine transform model. Then the predicted affinity (affinity*), watershed (watershed*) and agglomeration (seg*) were conducted based on the realigned chunk. The final segmentation (seg*) was inversely transformed back to the original space (seg). Affinity maps, 2D stacked watershed segmentations and agglomerated segmentations on x-y plane (top row) and x-z plane (bottom row). (**B**) Rendering example of the pair match process. Neurons in left and right chunk are shown in different color, and the overlapping area in mixed color. If two neuron objects’ Intersection over Union score exceeds the pre-defined threshold, they are merged as one neuron in global. Inset: example of pair match of axons. Scale bar, 5 μm. (**C**) An example of inter-chunk mergence which expands 29 chunks.

**Fig. S7.**
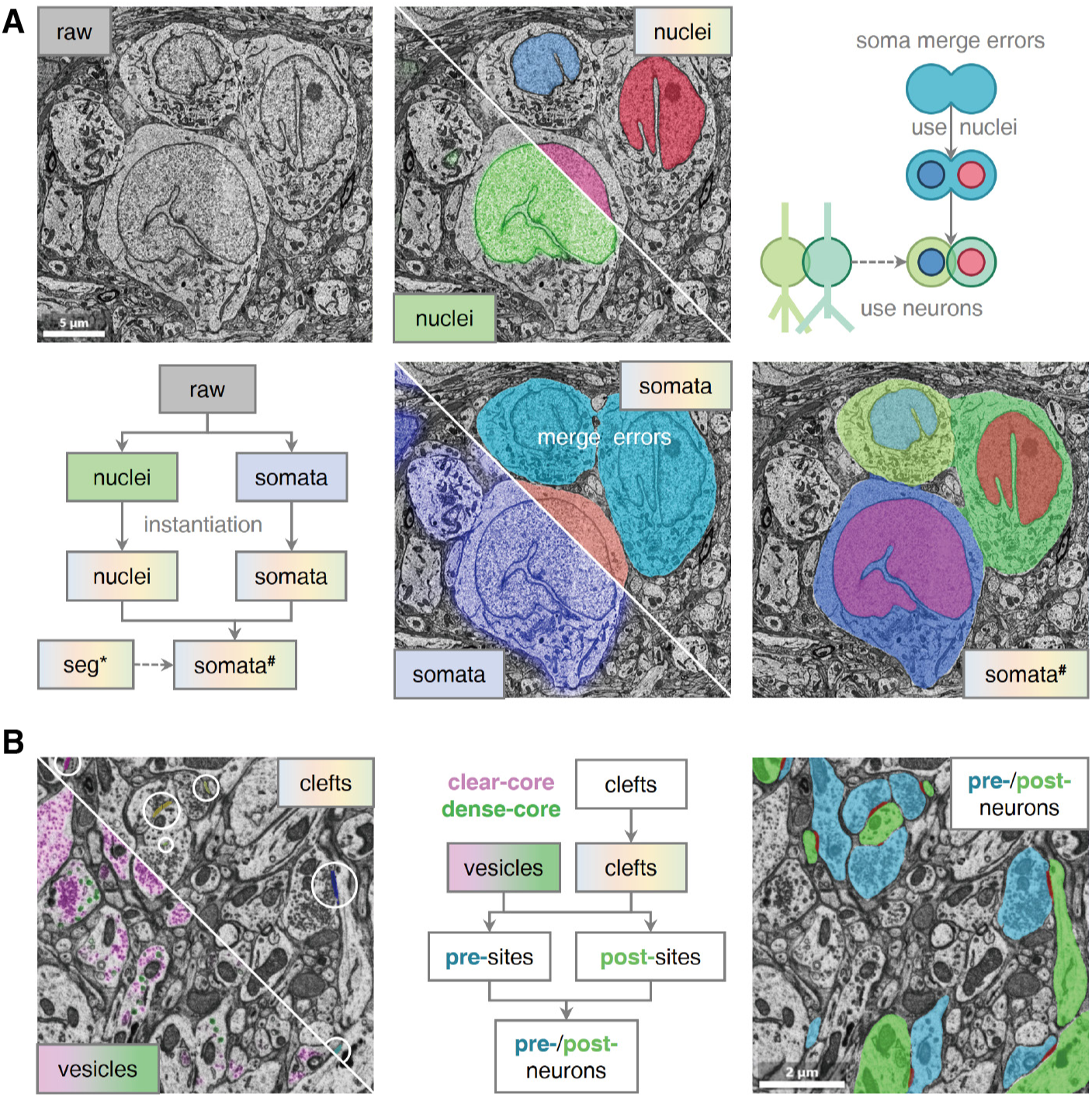
Pipeline for organelle segmentation. (**A**) Segmentation of nucleus and soma. Nuclei are detected at 40 × 40 × 40 nm^3^ resolution by a 3D CNN predicting the probability maps (light green, bottom left), followed by a connected component labeling (multicolor, top right) algorithm based on inter-slice overlap. Somata are detected at 80 × 80 × 40 nm^3^ resolution with a similar strategy, followed by a postprocessing step of correcting the merge errors based on nuclei. Scale bar, 5 μm. (**B**) Segmentation of vesicle, synapse and synaptic connectivity detection. Small synaptic vesicle (purple, bottom left) and dense core vesicle (dark green, bottom left) are detected and classified using a semantic segmentation network. Synapses are first segmented by a 3D CNN predicting the probability maps of each voxel and then reconstructed (multicolor, top right) by a connected components labeling algorithm. To determine the synaptic connectivity, a 3D CNN was used to predict the masks of pre-synaptic neuron (blue) and the post-synaptic neuron (green) for each cleft. All above is done at the resolution of 10 × 10 × 40 nm^3^. Scale bar, 2 μm.

**Table S1.**
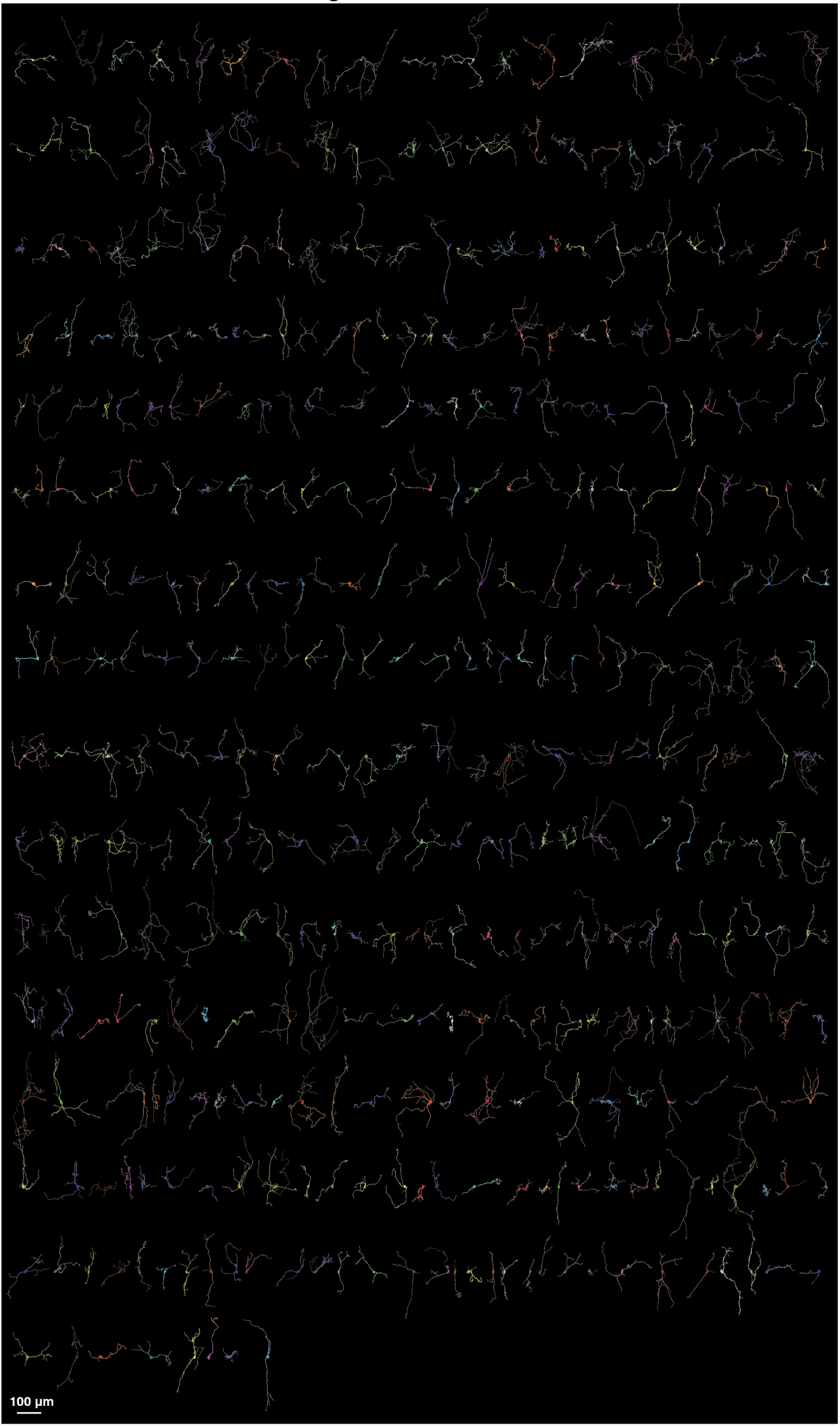
Neurons 1–391. Related to Fig. 2.

